# Establishment of chromatin accessibility by the conserved transcription factor Grainy head is developmentally regulated

**DOI:** 10.1101/798454

**Authors:** Markus Nevil, Tyler J. Gibson, Constantine Bartolutti, Anusha Iyengar, Melissa M Harrison

## Abstract

The dramatic changes in gene expression required for development necessitate the establishment of *cis-*regulatory modules defined by regions of accessible chromatin. Pioneer transcription factors have the unique property of binding closed chromatin and facilitating the establishment of these accessible regions. Nonetheless, much of how pioneer transcription factors coordinate changes in chromatin accessibility during development remains unknown. To determine whether pioneer-factor function is intrinsic to the protein or whether pioneering activity is developmentally modulated, we studied the highly conserved, essential transcription factor, Grainy head (Grh). Grh is expressed throughout *Drosophila* development and functions as a pioneer factor in the larvae. We demonstrated that Grh remains bound to condensed mitotic chromosomes, a property shared with other pioneer factors. By assaying chromatin accessibility in embryos lacking either maternal or zygotic Grh at three stages of development, we discovered that Grh is not required for chromatin accessibility in early embryogenesis, in contrast to its essential functions later in development. Our data reveal that the pioneering activity of Grh is temporally regulated and is likely influenced by additional factors expressed at a given developmental stage.

## Introduction

During metazoan embryonic development, cell-specific patterns of gene regulation are driven by complex transcriptional networks (Bonn and Furlong, 2008; Davidson, 2006). Transcriptional programs are orchestrated by *cis*-regulatory modules, such as enhancer and promoter elements (Bonn and Furlong, 2008; Davidson, 2006; Wittkopp and Kalay, 2012). These DNA elements provide modular platforms through which the input of multiple transcription factors integrate to output precise control of gene expression (Bonn and Furlong, 2008; Wilczyński and Furlong, 2010; Zinzen et al., 2009). While much progress has been made in defining specific features of *cis*-regulatory modules, less is known about the spatiotemporal mechanisms that establish individual modules during development.

The *cis-*regulatory modules of actively transcribed genes are located in “open” chromatin, or chromatin with low nucleosome occupancy and few high-order structures (Gross and Garrard, 1988; Kornberg, 1977; Kornberg and Lorch, 1999; Long et al., 2016). In contrast, “closed” chromatin has high-nucleosome density and extensive interactions between nucleosomes that present a barrier to transcription-factor binding (Kornberg, 1977; Luger et al., 2012; Tremethick, 2007). Accordingly, models of *in vivo* transcription-factor binding patterns are dramatically improved when chromatin accessibility is included along with *in vitro* binding affinities (Kaplan et al., 2011; Li et al., 2011). Thus, factors that influence chromatin structure help define the gene regulatory networks essential for development by establishing accessible *cis*-regulatory elements for transcription-factor binding.

A special class of transcription factors, termed pioneer factors, are able to overcome nucleosomal barriers to genome binding (Iwafuchi-Doi, 2019; Iwafuchi-Doi and Zaret, 2014; Soufi et al., 2012; Soufi et al., 2015; Zaret and Carroll, 2011). Pioneer factors bind closed chromatin to establish regions of accessibility, allowing other transcription factors to bind (Iwafuchi-Doi and Zaret, 2014; Soufi et al., 2012; Zaret and Carroll, 2011). Given their role in determining *cis-*regulatory modules, pioneer factors are hypothesized to act at the top of gene-regulatory networks (Iwafuchi-Doi and Zaret, 2014). A number of factors with pioneering activity have been identified that likely establish transcriptional networks required for cell identity: FOXA1 in mammalian liver stem cells; SOX2, OCT4, and KLF4 in induced pluripotent stem cells; Grainy head (Grh) in *Drosophila melanogaster* eye imaginal discs; and Zelda (Zld) in the early *Drosophila* embryo (Caravaca et al., 2013; Cirillo et al., 2002; Foo et al., 2014; Harrison et al., 2011; Jacobs et al., 2018; Schulz et al., 2015; Soufi et al., 2012; Soufi et al., 2015; Sun et al., 2015). However, it remains unclear whether these pioneering factors define accessible chromatin at multiple stages of development or whether their pioneering activity is limited to specific times in development or distinct tissues.

The Grh family of transcription factors is essential in defining epithelial cell fate (Boglev et al., 2011; Bray and Kafatos, 1991; Bray et al., 1988; Bray et al., 1989; Hemphälä et al., 2003; Nevil et al., 2017; Rifat et al., 2010; Traylor-Knowles et al., 2010; Uv et al., 1994; Uv et al., 1997; Wang and Samakovlis, 2012; Wilanowski et al., 2002; Yao et al., 2017). Grh proteins are highly conserved, with members found in all metazoans examined to date and with more distant relatives found in fungi (Paré et al., 2012; Traylor-Knowles et al., 2010; Venkatesan et al., 2003; Wang and Samakovlis, 2012; Wilanowski et al., 2002). Extensive work in *Drosophila* and mammalian cell culture has demonstrated that Grh acts to define an epithelial gene regulatory network during metazoan development (Boglev et al., 2011; Bray and Kafatos, 1991; Chen et al., 2016; Gao et al., 2013; Hemphälä et al., 2003; Nevil et al., 2017; Nishino et al., 2017; Senga et al., 2012; Wang and Samakovlis, 2012; Yao et al., 2017). More recently, Grh has been shown to have pioneer-factor activity, as it is required to maintain chromatin accessibility at enhancers in *Drosophila* eye imaginal discs and can poise these enhancers for transcriptional activation (Jacobs et al., 2018). This pioneering function is shared with mammalian GRHL proteins, which are required for defining enhancers in breast cancer cell lines and as cells exit from naïve pluripotency (Chen et al., 2018; Jacobs et al., 2018).

The fact that Grh is expressed throughout *Drosophila* development allows us to ask whether its pioneering activity is required at multiple developmental stages. By analyzing chromatin accessibility in embryos lacking either maternal or zygotic *grh*, we demonstrate that Grh is not required to define accessible *cis*-regulatory modules in the early embryo. This contrasts with its essential pioneering function in defining chromatin accessibility in the eye imaginal disc. By mutating an individual Grh-bound locus we show that additional factors may be able to compensate for loss of Grh even in the larvae. Thus, by analyzing the pioneering function of Grh at multiple developmental stages, it is evident that the role of this factor in defining accessible chromatin is modulated and may depend on additional factors expressed in a given tissue or at a specific time in development.

## Results

### Grainy head remains bound to mitotic chromatin during embryogenesis

While many DNA-binding factors do not remain on mitotic chromosomes, some vertebrate pioneer factors are retained on chromatin through mitosis (Bellec et al., 2018; Caravaca et al., 2013; Festuccia et al., 2016; Festuccia et al., 2017; Iwafuchi-Doi, 2019; Kadauke et al., 2012; Liu et al., 2017). This “mitotic bookmarking” activity is hypothesized to allow pioneer factors to rapidly re-establish transcriptional programs after cell division (Caravaca et al., 2013; Iwafuchi-Doi, 2019; Iwafuchi-Doi and Zaret, 2014). Given the role of Grh as a pioneer factor in larval development and the fact that its chromatin occupancy is remarkably stable through *Drosophila melanogaster* development (Jacobs et al., 2018; Nevil et al., 2017; Potier et al., 2014), we wanted to test whether Grh remained associated with the compacted mitotic chromosomes.

To visualize Grh expression and localization in living embryos, we engineered an N-terminal superfolder Green Fluorescent Protein (sfGFP)-tag on the endogenous protein (Fig. S1A; Gratz et al., 2013; Hamm et al., 2017; Pédelacq et al., 2006). This strategy is predicted to label a majority of known Grh isoforms. While *grh* null mutants are homozygous lethal (Bray and Kafatos, 1991; Bray et al., 1989; Hemphälä et al., 2003; Uv et al., 1994; Uv et al., 1997), strains homozygous for the sfGFP-Grh-encoding allele were viable and fertile with no obvious mutant phenotype, demonstrating that the sfGFP tag does not interfere with essential Grh function. Additionally, sfGFP-Grh was nuclear and expressed strongly in the epidermal tissue of developing embryos and the imaginal discs of larvae, where endogenous Grh expressed (Fig. 1A, S1B; Attardi and Tjian, 1993; Bray et al., 1989; Hemphälä et al., 2003; Lee and Adler, 2004; Narasimha et al., 2008).

**Figure 1:**
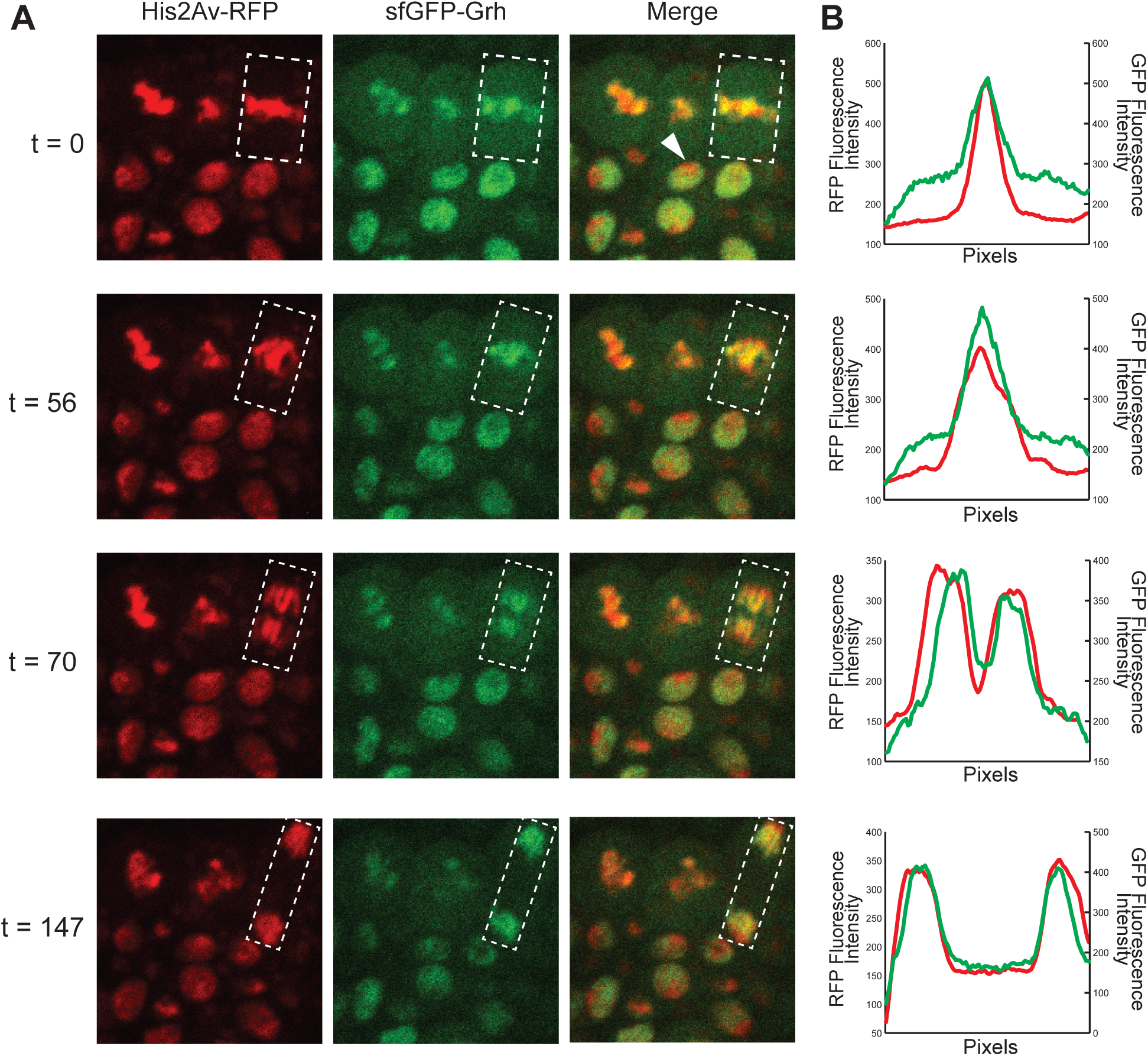
sfGFP-Grh remains on mitotic chromatin during gastrulation. (A) Cells of embryos expressing His2AV-RFP (chromatin) and sfGFP-Grh. White arrow indicates an interphase cell. White box indicates the region-of-interest where fluorescence intensity was measured. t, time in seconds from initial image. (see also Movie 1) (B) Relative fluorescence intensity for RFP (chromatin) and GFP (Grh), where the x axis is the total length of the region-of-interest.

In the early embryo, sfGFP-Grh was first expressed at detectable levels at gastrulation. During interphase, sfGFP-Grh was evident throughout the nucleus, except for a location we presume to be heterochromatin due to the strong fluorescent H2A (His2Av-RFP) signal (Fig. 1A t = 0, white arrow). sfGFP-Grh colocalized with the condensed chromosomes at metaphase, marked by His2Av-RFP, and this association with continued throughout mitosis (Fig. 1A, Movie 1). Measuring the relative fluorescence intensity across the nuclei of one mitotic event confirmed the strong correlation between Grh localization and mitotic chromatin (Fig. 1B). Together these results demonstrate that the pioneer factor Grh remains associated with compacted mitotic chromatin, a feature shared with other pioneer factors.

### Maternally supplied Grainy head is not essential for establishing chromatin accessibility in the early embryo

Grh plays an essential role in defining chromatin accessibility in the larval eye imaginal disc (Jacobs et al., 2018; Potier et al., 2014). We therefore sought to determine whether Grh influenced chromatin accessibility in the early embryo. *grh* is maternally deposited as an mRNA, and previous work demonstrated that this maternally encoded Grh binds to chromatin and is required for normal gene expression (Garcia and Stathopoulos, 2011; Harrison et al., 2010; Huang et al., 1995; Liaw et al., 1995; Nevil et al., 2017). We generated embryos depleted of maternally provided *grh* using the FLP/FRT system to generate mitotic clones of the *grh*^*B37*^ null mutant (Fig. S2A; Bray and Kafatos, 1991; Chou and Perrimon, 1996; Harrison et al., 2010; Nevil et al., 2017). Stage 5 embryos lacking maternal *grh* were identified by the absence of GFP fluorescence, and single embryos were collected in triplicate alongside wild-type sibling controls. Chromatin accessibility was measured with the assay for transposase-accessible chromatin using sequencing (ATAC-seq) (Buenrostro et al., 2013). We identified thousands of accessible regions at stage 5 in both wild-type and maternally depleted embryos. These largely overlapped with previously published data from the early embryo, confirming that our assay successfully identied accessible chromatin (Fig. S2B; Blythe and Wieschaus, 2016). Unexpectedly, there were no significant differences in chromatin accessibility between the maternal depletion (*grh*^*M-*^) and wild-type controls (*grh*^*M+*^) (Fig. 2, all q-values > 0.9 indicating no reproducible differential accessibility). Thus, although maternally encoded Grh is present and required for gene expression (Harrison et al., 2010; Huang et al., 1995; Nevil et al., 2017), it is not essential for establishing chromatin accessibility at stage 5.

**Figure 2.**
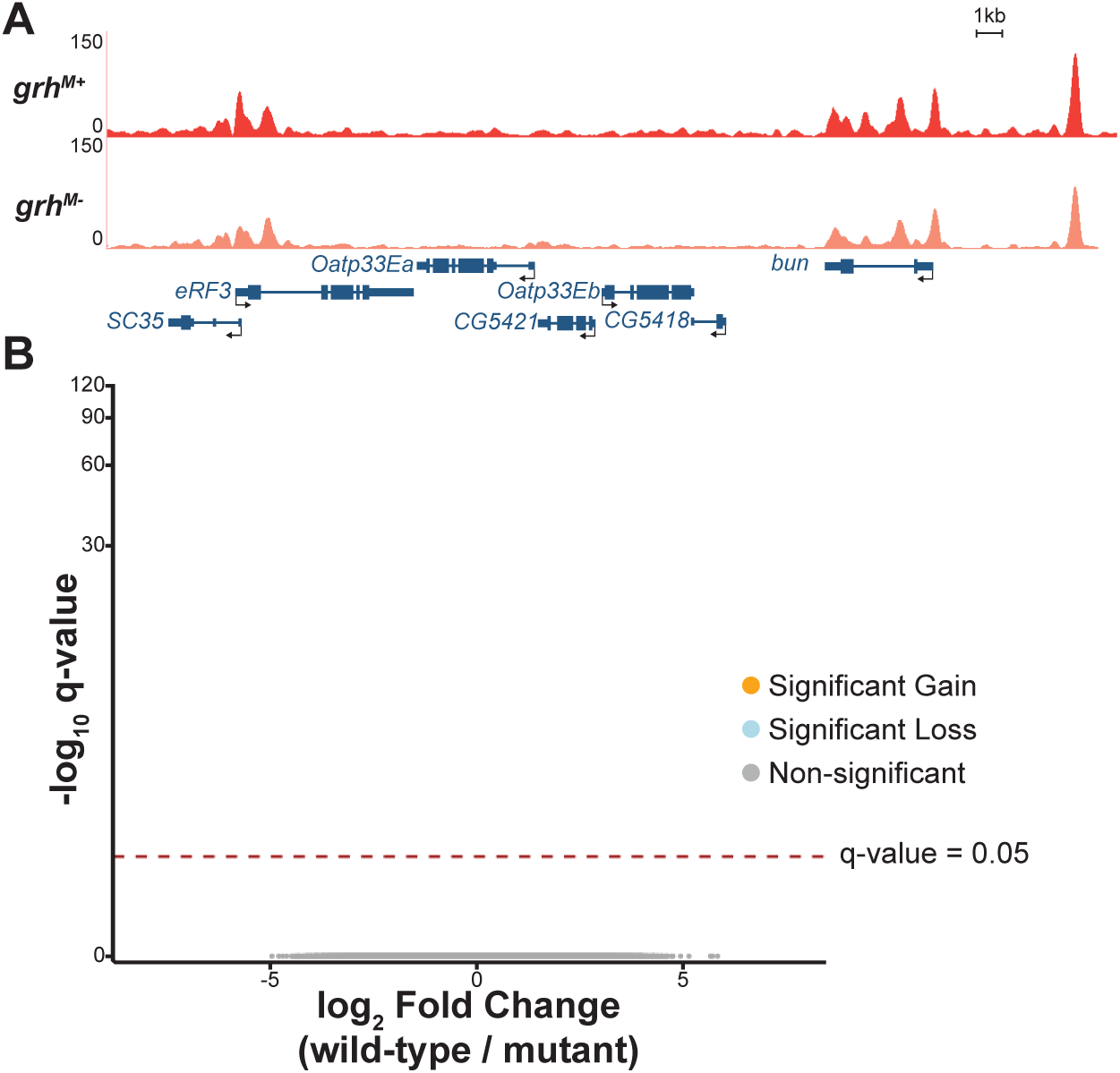
Maternally encoded Grh is not required for chromatin accessibility. (A) UCSC genome browser tracks of a representative locus showing single replicates of stage 5 wild-type control (*grh*^*M+*^) and maternal depletion (*grh*^*M-*^) ATAC-seq. (B) Volcano plots of all accessible regions in comparisons between the *grh* maternal depletion and heterozygous siblings. Significance of change in accessibility reported by -log_10_ q-value on the y-axis, and magnitude of change by log_2_ fold change on the x-axis. Regions that significantly gain (orange) or lose (blue) accessibility are defined as those with a q-value < 0.05. Non-significant changes are those with a q-value > 0.05 (gray).

### Widespread changes in chromatin accessibility accompany gastrulation

In addition to low levels of maternally encoded Grh, *grh* is also robustly expressed as the zygotic genome is activated during stage 5 (Sandler and Stathopoulos, 2016). While zygotic *grh* is ultimately required for embryogenesis, maternally encoded Grh is not required for survival (Harrison et al., 2010). To better understand the role of the essential zygotically encoded Grh, we determined the changes in the chromatin landscape that accompany the robust activation of *grh* expression as the zygotic genome is activated. We assayed chromatin accessibility in wild-type embryos at stage 6, immediately following the expression of zygotic *grh* (Fig. 3A). By comparing these data with our data from stage 5 embryos, we identified thousands of sites that gained or lost accessibility during gastrulation (Fig. 3B-E). These dramatic changes in accessibility did not require maternally supplied *grh* as we identified no significant differences in accessibility between stage 6 embryos depleted for maternally encoded Grh and their wild-type siblings (Fig. S3A). Hierarchical clustering of sample distances for stage 5 and stage 6 ATAC-seq datasets was driven more by time point than by genotype (Fig. S3B). These data suggest that developmental stage, and not loss of maternal *grh*, is the major driver altering the chromatin accessibility of these embryos.

**Figure 3.**
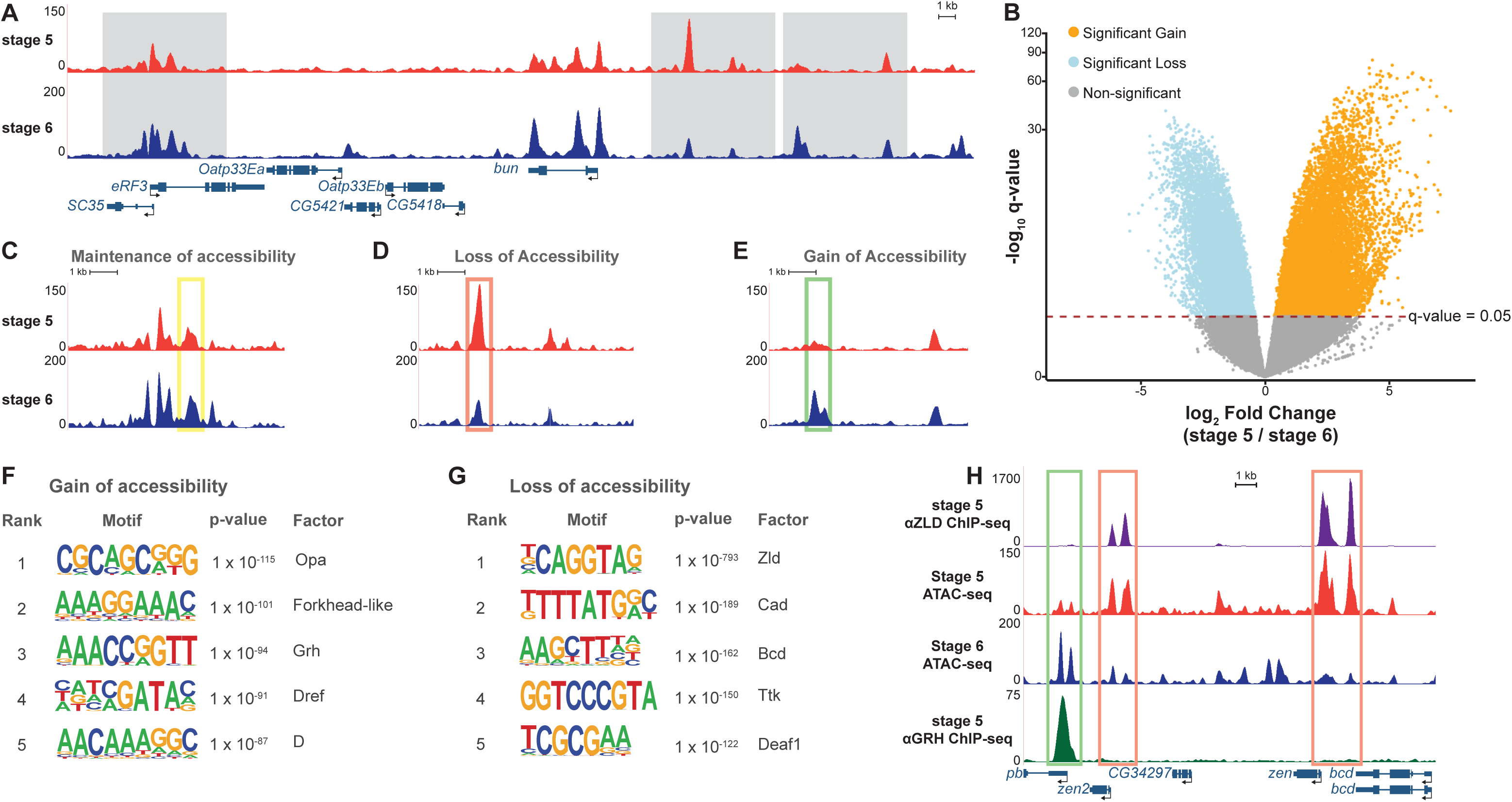
Widespread changes in chromatin accessibility occur during gastrulation. (A) UCSC genome browser track of a representative locus showing single replicates of stage 5 and stage 6 wild type (*grh*^*M+*^) ATAC-seq. (B) Volcano plot of all accessible regions identified in comparisons between stage 5 and stage 6. Significance of change in accessibility reported by -log_10_(q-value) on the y-axis, and magnitude of change by log_2_(fold change) on the x-axis. Regions that significantly gain (orange) or lose (blue) accessibility are defined as those with a q-value < 0.05. Non-significant changes are those with a q-value > 0.05 (gray). (C-E) Examples of accessible regions that maintain (C), lose (D), and gain (E) chromatin accessibility between stage 5 and stage 6 embryos. (F) Top 5 motifs identified in *de novo* motif enrichment of regions that gain chromatin accessibility at gastrulation. (G) Top 5 motifs identified in *de novo* motif enrichment of regions that lose chromatin accessibility at gastrulation. (H) UCSC Genome Browser tracks of ATAC-seq from stage 5 and stage 6 embryos with ChIP-seq data for Zelda (Zld) (Harrison et al., 2011) and Grainy head (Grh) (Nevil et al., 2017). Green box indicates regions with a significant gain in accessibility. Red boxes indicate regions with significant losses in accessibility.

To identify potential drivers of the widespread changes in chromatin accessibility during gastrulation, we performed *de novo* motif discovery for regions that gained and lost accessibility during gastrulation. Motifs for factors that are suggested to have roles as pioneer transcription factors were highly enriched in sites that gain accessibility during gastrulation (Fig. 3). Among them were Forkhead-like and Dichaete (D) whose orthologs in mammals (FOXA1 and SOX2, respectively) are known to have pioneering functions (Soufi et al., 2012; Zaret and Carroll, 2011). Additionally, the canonical Grh-binding motif was enriched at these sites that gain accessibility (Fig. 3F, p-value = 1e10^−94^). These results suggest that Forkhead-like, D, Grh, and other factors like Dref and Odd-paired (Opa) may have active roles in facilitating chromatin accessibility at gastrulation. This contrasts with regions that lost accessibility. The binding motif for the zygotic genome activator Zelda (Zld) was highly enriched in the sequences underlying regions that decreased in accessibility at gastrulation (Fig. 3G, p-value = 1e10^−793^; Liang et al., 2008). Thus, while Zld is a pioneer factor that functions to establish or maintain chromatin accessibility at a subset of loci in the stage 5 embryo, the enrichment of Zld-binding sites in areas with decreased accessibility suggests that other factors likely influence chromatin accessibility following genome activation (Foo et al., 2014; McDaniel et al., 2019; Schulz et al., 2015; Sun et al., 2015).

We have previously identified both Zld and Grh-binding sites at stage 5 using chromatin immunoprecipitation coupled with high-throughput sequencing (ChIP-seq) (Harrison et al., 2011; Nevil et al., 2017). Comparing our ChIP-seq and ATAC-seq data sets showed that Zld binding is enriched in regions that decrease in chromatin accessibility, while Grh binding is enriched in regions that gain chromatin accessibility (Fig. 3H, S3C), consistent with the binding predicted by motif searches. Thus, both motif enrichment and transcription-factor binding profiles support a role for these pioneering proteins in determining developmentally regulated chromatin accessibility.

### Zygotically expressed Grainy head is not required for chromatin accessibility at gastrulation

Because both the Grh-binding motif and Grh occupancy were enriched at regions that gained chromatin accessibility between stage 5 and stage 6, we tested whether zygotically encoded Grh was essential for driving these changes in chromatin architecture. We collected mutant (*grh*^*B37/B37*^) and wild-type (*grh*^*B37/+*^) stage 6 embryos in triplicate and measured the chromatin accessibility using single embryo ATAC-seq (Fig. S4A, B). While we identified hundreds of sites that lose accessibility in the *grh* mutant background, these sites were false positives. 90.6% of the regions that changed in accessibility were located on the same chromosome as *grh*, chromosome 2 (Fig. 4A, B). Because the *grh*^*B37*^ allele is maintained over a balancer chromosome this *grh*^*B37*^-containing chromosome has likely accumulated mutations (Sharp and Agrawal, 2012; Sharp and Agrawal, 2017), which prevented proper read mapping, artificially lowering read counts and resulting in apparent losses of accessibility (Fig. S3C). Thus, although both the Grh motif and Grh binding were enriched in regions that gained in accessibility during gastrulation, loss of zygotic *grh* alone is not sufficient to significantly alter chromatin accessibility in stage 6 embryos.

**Figure 4.**
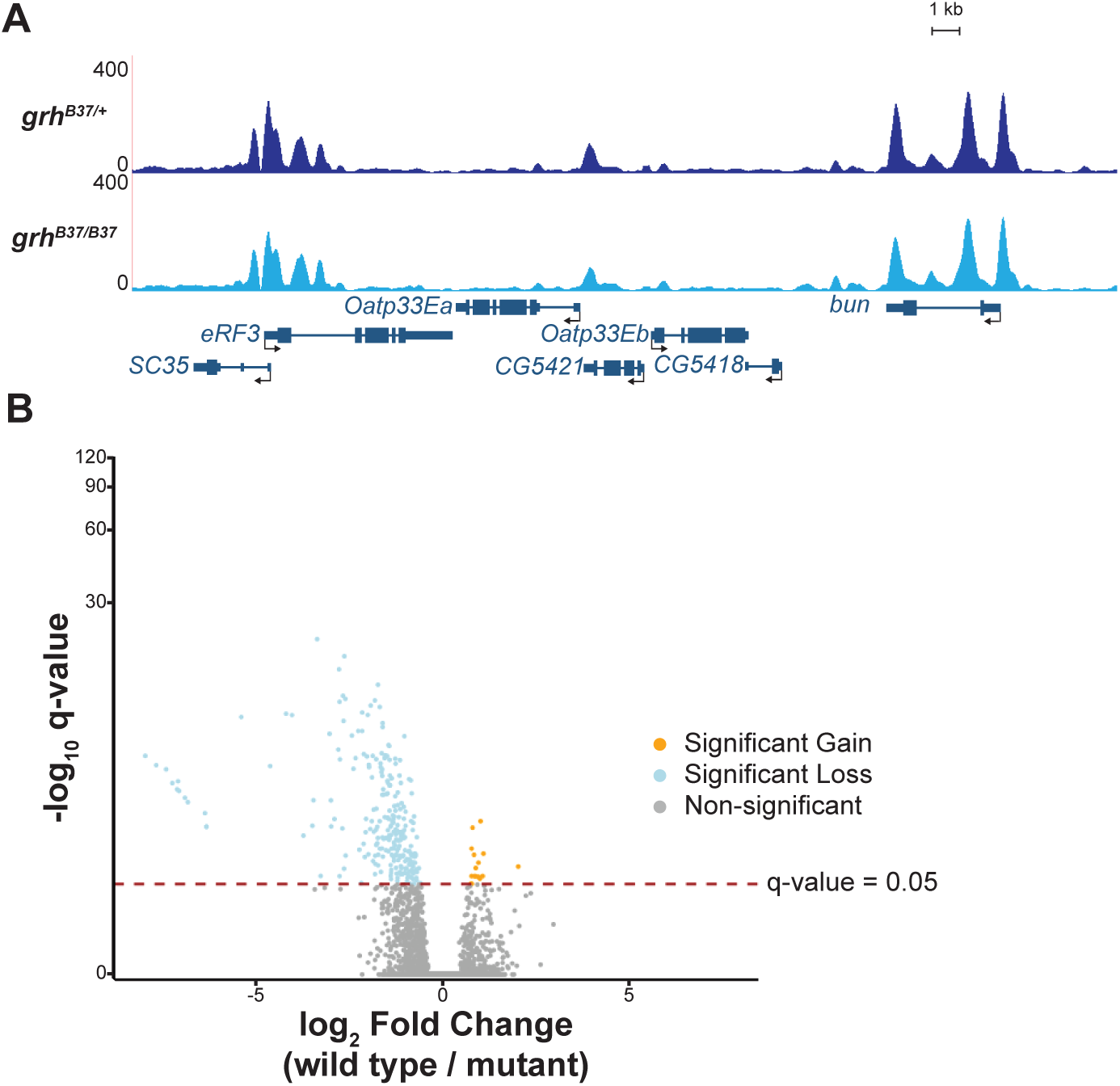
Zygotic Grh is not required for chromatin accessibility at gastrulation. (A) UCSC genome browser tracks of single stage 6 embryo ATAC-seq data from either a *grh* null mutant (*grh*^*B37/B37*^) or wild-type sibling control (*grh*^*B37/+*^). (B) Volcano plot showing all accessible regions identified. Significance of change in accessibility reported by -log_10_(q-value) on the y-axis, and magnitude of change by log_2_(fold change) on the x-axis. Regions that significantly gain (orange) or lose (blue) accessibility are defined as those with a q-value < 0.05. Non-significant changes are those with a q-value > 0.05 (gray) (see also Fig. S4).

### Grainy head maintains chromatin accessibility during late stages of embryogenesis

Our data showed that contrary to our expectations, early chromatin accessibility is not dependent upon either maternally encoded or zygotically expressed Grh. However, since Grh is essential for chromatin accessibility in larval imaginal discs (Jacobs et al., 2018), we tested whether later in embryonic development Grh may be important for maintaining or establishing accessible regions in cells as they differentiate. To test this, we depleted zygotic Grh and performed bulk ATAC-seq on homozygous *grh*^*B37*^ mutant embryos collected 11-12 hours after egg laying (AEL, stage 14-15) and their heterozygous sibling controls. Embryos at this stage are comprised of numerous different tissue types, many of which do not express Grh. Despite this heterogeneity, we identified 92 regions with significant losses in chromatin accessibility in the absence of Grh (Fig. 5A, B). Our ability to identify these changes in accessibility within whole embryos suggested that the 92 identified regions may be especially dependent on Grh for accessibility. Thus, to determine if Grh was more broadly required for accessibility we examined whether there was a general decrease in accessibility at all sites bound by Grh when Grh was absent. We identified a dependence on Grh for accessibility when we analyzed 5 kb windows surrounding regions normally bound by Grh (Fig. 5C; Nevil et al., 2017). To quantify this observation, we tested for changes in accessibility in windows around Grh-binding sites. We normalized read depth in 50 bp windowed regions around all Grh-peak centers and measured the loss of accessibility as the log_2_ ratio of mutant:wild-type normalized read depth. We noted the greatest difference between mutant and wild type in accessibility is at Grh-peak centers (Fig. 5D, compare “Genomic” to “0-50bp”). Furthermore, this decrease in accessibility as compared to wild type was alleviated as distance from the binding-site center increased (Fig. 5D, compare “0-50 bp” to “1000-1050 bp”). To test whether this dependence on Grh for accessibility was specific to distinct genomic regions, we annotated and separated the genome into promoters, 1-3kb upstream, exons, introns, untranslated regions (UTRs), and intergenic regions. We then identified the changes in accessibility at these regions in the presence or absence of Grh. Independent of genomic feature, the loss of Grh resulted in decreased chromatin accessibility at regions normally bound by Grh (Fig. S5). Together, we identified a role for Grh in establishing or maintaining accessibility during the late stages of embryogenesis.

**Figure 5.**
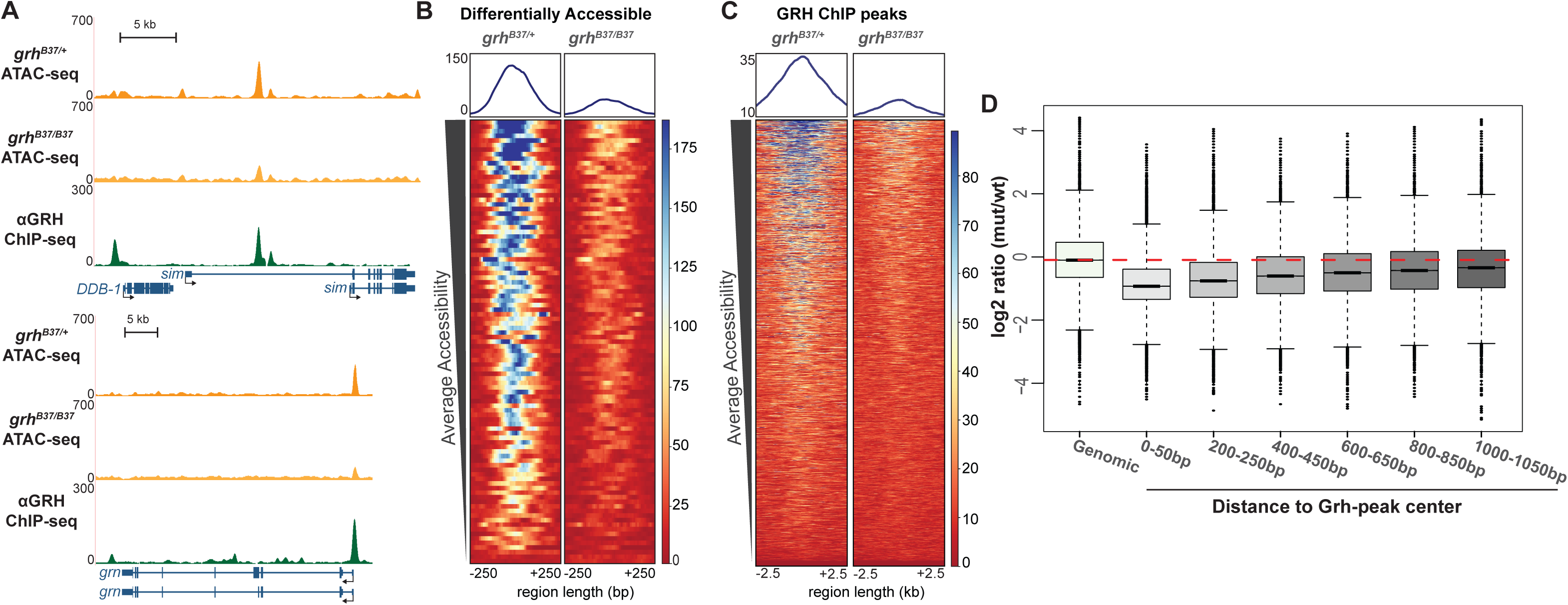
Grh is required for chromatin accessibility 11-12 hour after egg laying. (A) UCSC genome browser tracks of ATAC-seq from 11-12hr AEL (hours after egg laying) for *grh-*null mutant embryos (*grh*^*B37/B37*^), stage-matched, wild-type sibling control (*grh*^*B37/+*^), and stage-matched Grh ChIP-seq (Nevil et al., 2017). (B) Heatmaps of regions differentially accessible between the Grh mutant and wild-type control. Color scale indicates relative height of ATAC-seq, i.e. accessibility. Heatmaps centered on ATAC-seq peak summits. (C) Heatmaps of ATAC-seq data from *grh*-mutant and wild-type control embryos for all Grh-bound regions as identified by ChIP-seq. Color scale indicates relative height of ATAC-seq, i.e. accessibility. Heatmaps are centered on Grh ChIP-seq peak summits. (D) Box plots of log_2_ ratios of ATAC-seq signal between *grh*-mutant and wild-type embryos, at windows of increasing distance around Grh ChIP-seq peaks. Red-dotted line indicates average signal of random genomic windows.

### Loss of Grainy head at a single locus does not significantly alter chromatin accessibility

Removing Grh in the embryo also disrupts the expression of additional transcription or developmental factors controlled by the Grh gene-regulatory network (Nevil et al., 2017). To determine the direct effects of removing the ability of Grh to bind to the genome, we mutated a single Grh-binding motif using Cas9-mediated genome editing and assayed chromatin accessibility using formaldehyde-assisted isolation of regulatory elements (FAIRE) coupled with qPCR. For this purpose, we identified a single canonical Grh-binding motif in the promoter of the gene *ladybird late* (*lbl*). Grh is bound to this locus throughout embryogenesis and in larval wing discs (Fig. 6; Nevil et al., 2017). This site gained accessibility during gastrulation and loses accessibility in the absence of zygotic Grh at 11-12 hours after egg laying (AEL) (Fig. 6A).

**Figure 6.**
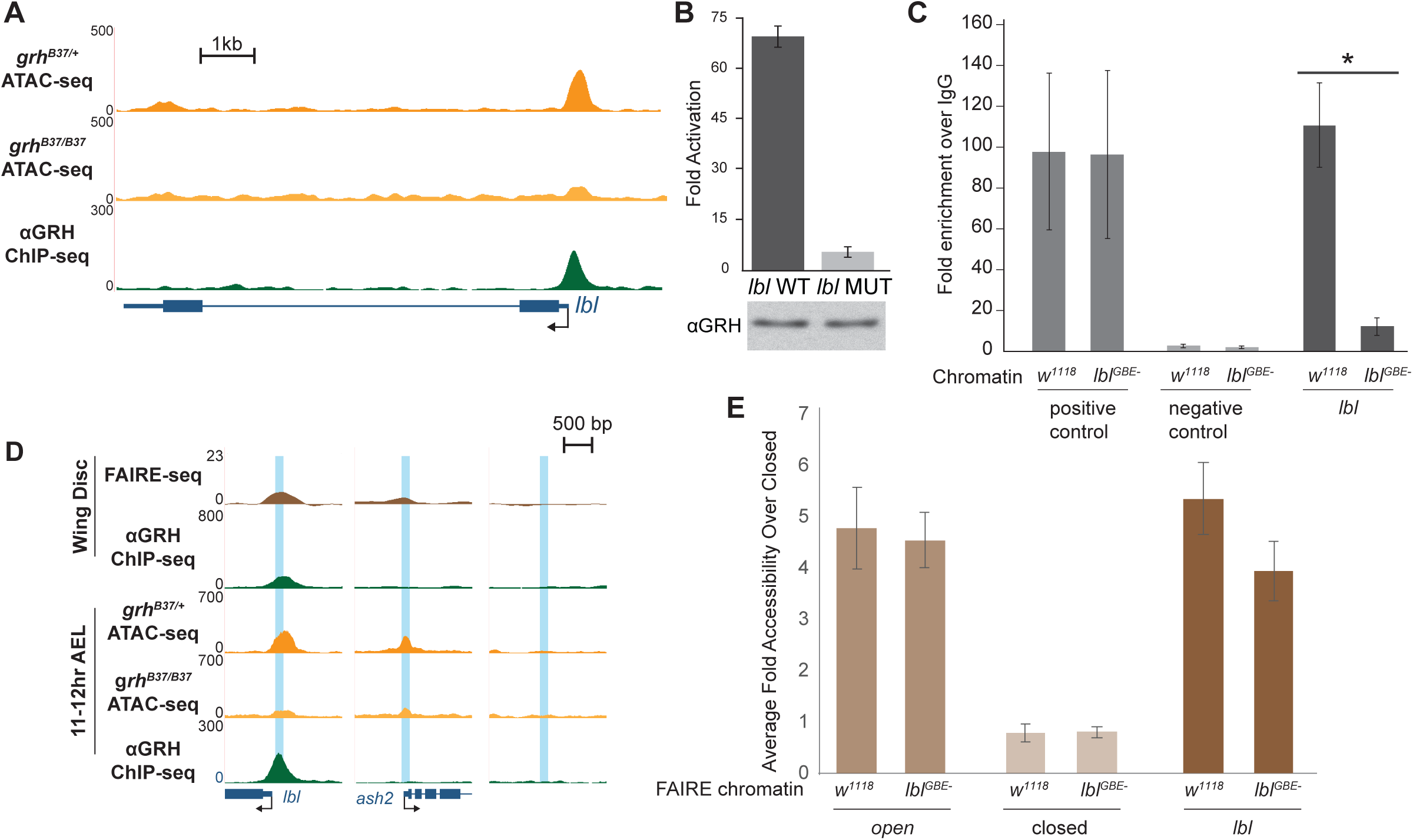
Grh is not required for local accessibility at the *ladybird late* promoter. (A) UCSC genome browser tracks of 11-12hr AEL ATAC- and ChIP-seq data at the *lbl* locus for the genotypes indicated. (B) Fold activation upon Grh expression of reporters driven by either a wild-type *lbl* promoter (*lbl* WT) or the *lbl* promoter with the Grh-binding site mutated (*lbl* MUT) transiently transfected into S2 cells. Error bars indicate the standard deviation (C) ChIP-qPCR using an anti-Grh antibody performed with chromatin from either wild-type (*w*^*1118*^) embryos or embryos in which the single Grh-binding site in the *lbl* promoter had been disrupted (*lbl*^*GBE-*^). Positive control, Grh-bound locus (*Slo*). Negative control, intergenic region. *lbl*, region corresponding to Grh-binding site in the *lbl* promoter. Error bars indicate the standard deviation, * indicates p-value < 0.05 calculated using a t-test, n = 3. (D) UCSC genome browser tracks of ATAC-, ChIP-(Nevil et al., 2017), and FAIRE-seq data (McKay and Lieb, 2013) from of 11-12hr AEL embryos and larval imaginal wing discs. Blue boxes indicate target regions assayed by FAIRE. (E) FAIRE-qPCR results from *lbl* promoter mutant (*lbl*^*GBE-*^) and wild-type (*w*^*1118*^) chromatin at an open region (*ash2*), closed region, and the *lbl* promoter. Error bars indicate the standard deviation, n = 3.

To confirm the necessity of the identified Grh-binding motif to recruit Grh, we transiently transfected S2 cells with a luciferase reporter driven by the *lbl* promoter with either a wild-type or mutated Grh motif and assayed luciferase levels in the presence or absence of Grh. Grh activated expression of the wild-type reporter, but not the reporter with the mutated binding motif (Fig. 6B). We therefore used Cas9-mediated mutagenesis to mutate the canonical Grh motif to an EcoRI-recognition site at the endogenous *lbl* locus (Fig. S6). To our knowledge this only disrupted the Grh motif and no additional promoter elements. The resultant flies were homozygous viable and fertile. ChIP-qPCR confirmed the loss of Grh binding to the mutant *lbl* promoter (*lbl*^*GBE-*^) and no detectable changes in binding to a control target regions (Fig. 6C). To assay for any changes in chromatin accessibility, we performed FAIRE-qPCR using 3^rd^ instar larval wing discs. Wing discs are a useful tissue for testing the role of Grh as the *lbl* promoter is accessible (McKay and Lieb, 2013), Grh is bound to the promoter (Nevil et al., 2017), and Grh is strongly expressed throughout the tissue (Fig. 6D, note imaginal disc ChIP- and FAIRE-seq; Lee and Adler, 2004; Nevil et al., 2017; Uv et al., 1997). FAIRE-qPCR of the mutated *lbl* promoter revealed a small, but statistically insignificant change in accessibility at the *lbl* promoter (Fig. 6E). Therefore, although Grh is required for chromatin accessibility at some larval eye disc enhancers (Jacobs et al., 2018), it is not required at all *cis-*regulatory modules.

## Discussion

We identified that the pioneering activity of *Drosophila* Grh is developmentally regulated. Previous work had demonstrated that Grh was both necessary and sufficient for chromatin accessibility in the larval eye disc and that GRHL2 similarly had a pioneering function in mammalian cell culture (Chen et al., 2018; Jacobs et al., 2018). Here we tested both the requirement for maternally and zygotically encoded Grh in determining regions of open chromatin in embryo. We demonstrated that maternally encoded Grh is not required for chromatin accessibility at stage 5, and zygotically encoded Grh is not required at stage 6, when zygotic *grh* is normally expressed. Nonetheless, Grh motifs are one of several sequence motifs enriched at sites that become accessible during gastrulation. In contrast to gastrulating embryos, we established that Grh activity is important for determining chromatin accessibility later in embryonic development. This is also the developmental time point at which Grh is essential for viability. Despite this role in determining chromatin accessibility in the late embryo and larvae, the loss of Grh binding at a single locus in larval tissues did not decrease chromatin accessibility at this site. Thus, we propose that the pioneering activity of Grh is not required at all stages of development nor at all Grh-bound *cis-* regulatory elements. Instead, in these tissues or at these loci other factors may compensate for the loss of Grh.

While pioneer factors are defined by their ability to establish gene regulatory networks by binding to *cis-*regulatory modules and promoting chromatin accessibility, recent evidence suggests that pioneer factors vary in their capacity to accomplish this task. FOXA1, which is known to displace nucleosomes and to bind mitotic chromatin (Caravaca et al., 2013; Cirillo et al., 2002; Zaret and Carroll, 2011), is redirected to previously unoccupied sites upon activation of the TNFα pathway (Franco et al., 2015). Similarly, OCT4 binding to the genome is dynamic and is modulated by a cohort of transcription factors, including OTX2, to compete with nucleosomes at enhancers, (Buecker et al., 2014), and SOX2 requires PARP-1 to reshape nucleosomal DNA to access 26% of its sites *in vivo* (Liu and Kraus, 2017). Thus, the pioneering roles of these transcription factors are regulated in a tissue-specific or temporal manner. By assaying the conserved, essential transcription factor, Grh, at multiple stages of development, we demonstrated that the role of Grh in determining *cis*-regulatory modules depends on developmental stage. While Grh functions as a pioneer factor at a subset of enhancers in *Drosophila* eye imaginal discs and in mammalian epiblast stem cells (Chen et al., 2018; Jacobs et al., 2018), our data show that this activity is not required to establish or maintain chromatin accessibility in the early embryo. Together these data suggest that there is context specificity to Grh pioneering activity and support a model in which pioneer-factor activity is regulated by additional factors whose expression is variable across development.

The conditions that lead to context-specific Grh-pioneering activity remain unknown, but the combinatorial action of other transcription factors could provide temporal robustness to chromatin remodeling. By examining changes in chromatin accessibility during gastrulation, we have identified factors that may function to define *cis*-regulatory regions at this stage of development. Among the top motifs associated with gains in accessibility during gastrulation are binding motifs for the products of *odd-paired* (*opa*), *forkhead-like*, and *Dichaete* (*D*). The mammalian orthologs of these *Drosophila* transcription factors mark *cis-*regulatory regions. Zic2, the mammalian ortholog of Opa, occupies enhancers prior to OCT4 binding and thus differentiation, suggesting that Zic2 has a role in marking *cis-*regulatory regions (Luo et al., 2015). Forkhead-like factors in *Drosophila* are the homologs of FOXA1, a pioneer factor that actively displaces nucleosomes (Cirillo et al., 2002). *Dichaete* is a *Drosophila* member of the Group B Sox transcription factors (Aleksic et al., 2013; Crémazy et al., 2001; Russell et al., 1996). In mammals, the Group B Sox protein SOX2 is known to be a pioneer factor essential for development (Iwafuchi-Doi and Zaret, 2014; Soufi et al., 2012). In *Drosophila*, both *opa* and *D* are up-regulated upon zygotic genome activation and both are required for embryonic development (Benedyk et al., 1994; Nambu and Nambu, 1996). Opa is required for temporally regulated changes in expression of multiple pair-rule genes and functions both directly to induce spatiotemporal changes in expression and to modify the role of additional factors (Clark and Akam, 2016). D binds to core promoters and enhancers in the embryo and is required for proper gene expression of thousands of genes (Aleksic et al., 2013). Thus, Opa, Grh, and D are all factors that are broadly expressed at the transition to gastrulation and are required for proper gene expression. Together with our data demonstrating an enrichment for their binding motifs at regions of chromatin that become accessible at gastrulation, this suggests that these factors collaborate to determine the *cis*-regulatory regions. Furthermore, D and Opa may compensate for the loss of Grh at a subset of co-occupied regions at this time in development (Figure S7). However, later in development Grh is required broadly to establish chromatin accessibility (Jacobs et al., 2018), suggesting these and other factors can no longer compensate. Indeed in the larval brain, D, Opa and Grh constitute a non-redundant temporal cascade regulating neuroblast fate in the larval brain (Abdusselamoglu et al., 2019). This supports a model by which the requirement for pioneer-factor activity in determining *cis*-regulatory regions is not strictly inherent to the protein, but is dependent on the developmental stage in which the protein is acting.

During mitosis, chromatin condensation leads to the removal of many transcription factors from their interphase binding sites, but recent studies have indicated that a subset of factors remain bound to mitotic chromatin (Raccaud and Suter, 2018; Raccaud et al., 2019). While not a unique property of pioneer transcription factors, mitotic chromatin occupancy is correlated with the ability of pioneer factors to bind nucleosomal DNA and may allow these factors to rapidly reestablish transcriptional networks following mitosis (Caravaca et al., 2013; Festuccia et al., 2017; Kadauke et al., 2012; Soufi et al., 2012). We showed that Grh binds to mitotic chromatin in the gastrulating embryo. However, at the same time in development Grh is not essential for defining regions of chromatin accessibility. Thus, our data separate the pioneering activity and mitotic-chromatin binding activities of Grh. While the mechanisms and consequences of the retention of Grh on mitotic chromosomes in the early embryo are unclear, this ability may be related to its surprisingly stable binding profiles during embryogenesis as assayed by ChIP-seq (Nevil et al., 2017).

We previously demonstrated that Grh binding is stable across days of development (Nevil et al., 2017). By contrast, here we show that Grh activity in defining *cis*-regulatory regions is regulated during development. Our analysis suggests a number of factors that may have pioneering roles at gastrulation that compensate for loss of Grh. Furthermore, our results demonstrate that Grh remains bound to chromatin during mitosis, but that this function is not directly related to its pioneering function. Together, our data support a model in which pioneering activity is not a static property of the protein but is rather regulated and context dependent.

## Materials and Methods

### Fly stocks

All fly stocks were raised on molasses food at 25°C. Germline mitotic clones were produced as described in (Harrison et al., 2010) using the heat shock inducible FLP-FRT system (Chou and Perrimon, 1996). For zygotic Grh depletions, the *grh*^*B37*^ null mutant (Bray and Kafatos, 1991) was balanced over CyO sChFP. Embryos selected at stage 5, stage 6, or 11-12hr AEL (after egg laying) were screened by the absence of red fluorescence (11-12hr AEL zygotic depletion), green fluorescence (stage 5 and stage 6 maternal depletions, Fig. S2A), or by PCR screening (stage 6 zygotic depletion, Fig. S3A). Embryos were precisely staged using a light microscope. Additional fly strains used: *sfGFP-Grh* (this study); *His2Av-RFP(III)* (Bloomington *Drosophila* Stock Center (BDSC) #23650), *w*^*1118*^;*CyO, P{Tub-PBac T}2/Sp;l(3)/TM6B* (BSC #8285), *lbl*^*GBE-*^ (this study), and *w*_*1118*_.

### CRISPR/Cas9-mediated generation of sfGFP-Grh and *lbl* mutant promoter

Cas9-mediated genome engineering as described in (Gratz et al., 2013; Hamm et al., 2017) was used to generate the N-terminal super folder Green Fluorescent Protein (sfGFP)-tagged Grh and mutant *lbl* promoter. The double-stranded DNA (dsDNA) donor was created using Gibson assembly (New England BioLabs) 1-kb homology arms flanking the sfGFP tag and Grh N-terminal open reading frame. Additionally, a 3xP3-DsRed cassette flanked by the long-terminal repeats of PiggyBac transposase was placed in the Grh 5’ UTR for selection. The guide RNA sequence (GCCAACTCCTAGGCGGCTGT) was cloned into pBSK under the U63 promoter using inverse PCR. Purified plasmid was injected into embryos of *w[1118]; PBac{y[+mDint2]=vas-Cas9}VK00027* (BDSC#51324) by BestGene Inc. Lines were screened for DsRed expression to verify integration. The entire 3xP3-DsRed cassette was cleanly removed using piggyBac transposase. Sanger sequencing of the entire region confirmed integration of the sfGFP tag without errors.

For production of a fly strain lacking the Grainy head-binding element (GBE) in the *lbl* promoter (*lbl*^*GBE-*^), a single-stranded oligodeoxynucleotide (ssODN) donor with mutations in the Grh motif was produced by Integrated DNA Technologies. A guide RNA site was selected that overlapped the Grh motif. The modification of the Grh motif (AACTGGTT) to an EcoRI site (AAGAATTC) in the ssODN was sufficient to disrupt the PAM site and seed sequence, in addition to providing a restriction cut site for screening. The guide RNA sequence (TTTGGGGCCTCCAAACTGGT) was cloned into pBSK under the U63 promoter using inverse PCR. The ssODN was injected into embryos of *w[1118]; PBac{y[+mDint2]=vas-Cas9}VK00037/CyO, P{w[+mC]=Tb[1]}Cpr[CyO-A]* (BDSC#56552) by BestGene Inc. Lines were screened using PCR and EcoRI digestion. The promoter was then sequenced to confirm mutation without errors.

### Imaging live embryos

Homozygous 0 - 2 hour old *sfGFP-Grh*; *His2AV-RFP* embryos were dechorionated in 50% bleach for 2 minutes and subsequently mounted in halocarbon oil. The living embryos were imaged on a Nikon A1R+ confocal at the University of Wisconsin-Madison Biochemistry Department Optical Core. Staging was performed on the microscope to identify gastrulating embryos. Mitosis was identified in cells with condensing chromatin (visualized with *His2AV-RFP*) and cytoplasmic sfGFP-Grh (released upon nuclear envelope degradation). Videos were acquired immediately upon entry into mitosis. FIJI (Schindelin et al., 2012) was used to measure the relative fluorescence intensity of the fluorophores.

### ATAC-seq and bioinformatic analysis

Maternal *grh* depletions were obtained as described above. Homozygous zygotic *grh* mutants (*grh*^*B37/B37*^) at stage 6 were identified after ATAC-seq library preparation by PCR followed Sanger sequencing and subsequently confirmed after high-throughput sequencing (Fig. S4A-B). Homozygous zygotic *grh* mutants at 11-12 hours after egg laying were identified by lack of fluorescence from a CyO sChFP balancer chromosome.

Single-embryo ATAC-seq was performed as described previously (Blythe and Wieschaus, 2016; Buenrostro et al., 2013). Briefly, a single dechorionated embryo was transferred to the detached cap of a 1.5 mL microcentrifuge tube containing 10 µL of ice-cold ATAC lysis buffer (10 mM Tris pH 7.5, 10 mM NaCl, 3 mM MgCl_2_, 0.1% NP-40). Under a dissecting microscope, a microcapillary tube was used to homogenize the embryo. The cap was placed into a 1.5 mL microcentrifuge tube containing an additional 40 µL of cold lysis buffer. Tubes were centrifuged for 10 minutes at 500 x g at 4°C. ATAC-seq library preparation was performed using the Illumina Nextera DNA library preparation kit. The supernatant was removed, and the resulting nuclear pellet was resuspended in 5 µL buffer TD and combined with 2.5 µL H_2_O and 2.5 µL tagment DNA enzyme. Tubes were placed at 37°C for 30 minutes and the resulting fragmented DNA was purified using the Qiagen Minelute Cleanup Kit, with elution performed in 10 µL of the provided elution buffer. Fragmented DNA was amplified and barcoded with 12 cycles of PCR using New England BioLabs NEBNext Hi-Fi 2x PCR master mix and indexed primers. Amplified libraries were purified using a 1.2x ratio of Axygen Axyprep Mag PCR Cleanup beads. Libraries were submitted for paired-end, 150 bp sequencing at Novogene Inc. using an Illumina HiSeq 4000.

Bulk embryo ATAC-seq was performed on embryos using an ATAC-seq protocol described in (Buenrostro et al., 2013). Briefly, seven staged and dechorionated embryos were transferred to a 1.5 mL microcentrifuge tube and homogenized in 40 µL ice-cold ATAC lysis buffer. Library preparation and clean-up continued as described above. A 1.8x ratio of AMPure XP beads (Beckman Coulter) were used to purify the amplified libraries. Libraries were submitted for single-end, 100 bp sequencing at the University of Wisconsin-Madison Biotechnology Center DNA Sequencing Facility using an Illumina HiSeq 2500.

Using FASTQC paired- and single-end reads were checked for quality (Simon Andrews). For the paired-end reads adapters, Tn5 transposon sequences, and low-quality bases were removed using Trimmomatic: java -jar trimmomatic-0.36.jar PE -trimlog trimlog.txt raw_reads_P1.fastq.gz raw_reads_P2.fastq.gz P1_trim.fastq.gz U1_trim.fastq.gz P2_trim.fastq.gz U2_trim.fastq.gz NexteraPE-PE.fa:2:30:10:8:true LEADING:3 TRAILING:3 SLIDINGWINDOW:4:22 MINLEN:1 (Bolger et al., 2014). Concordant, paired reads were mapped to the dm6 genome assembly (dos Santos et al., 2015) using `Bowtie 2: bowtie2 –dovetail -k 2 -p 4 -N 1 -R 3 -t – met-file metrics.txt -x dm6 −1 P1_trim.fastq.gz −2 P2_trim.fastq.gz` (Langmead and Salzberg, 2012). Unmapped, multiply aligning, mitochondrial, and scaffold reads were removed. Fragments greater than 101 bp in length were removed leaving fragments largely originating from open chromatin. MACS version 2 (Zhang et al., 2008) was used with default parameters to identify any regions of open chromatin in all data sets: `macs2 callpeak -q 0.05 –call-summits -f BAMPE -t replicate_1 replicate_2 replicate_3 -n output`. These accessible sites were then combined into a single data set and merged to create a master list of all sites assayed in 100 bp regions. Throughout Samtools was used to filter and convert file formats (Li et al., 2009). Read counts of each accessible site for all replicates was obtained using featureCounts: `featureCounts -F SAF -O -p -P -d 1 -D 100 -a accessible_peaks.saf -o count_table.txt [BAM file list]` (Liao et al., 2013). The resultant table was used in R (R Core Team, 2017) using the DESeq2 package (Love et al., 2014) to test for differential accessibility. Significant results are those with q-value < 0.05 to minimize false positives. Subsequent visualization of DEseq2 tables were done using the ggplot2 package (Wickham, 2009). Visualization of genomic data was achieved by generation of bigWig files (Kent et al., 2010) and display at the UCSC Genome Browser (*http://genome.ucsc.edu*) (Kent et al., 2002; Raney et al., 2014). Single-end libraries were analyzed as described above with the appropriate default single-end parameters for each tool. No *in silico* size selection was performed on single-end data, as fragment size is not captured by this sequencing method.

### Motif enrichment

To test for enrichment of motifs, a *de novo* motif search was done using Hypergeometirc Optimization of Motif Enrichment (HOMER) (Heinz et al., 2010). Sites of significant changes in accessibility (identified by DEseq2) were used as input. The program identified motifs enriched relative to the background. The background for increases in accessibility were the sites that decreased in accessibility and vis versa. The *de novo* motifs were matched to known motifs from the JASPAR (http://jaspar.genereg.net) and DMMPMM (http://autosome.ru/DMMPMM) databases by HOMER. Motifs are given a p-value indicating the confidence of the enrichment relative to the background sequences.

### Cell culture and luciferase assays

The promoter region (−162 bp to +397 bp) from *lbl* was cloned into pGL3-Basic (Promega) to drive expression of Firefly luciferase. The canonical Grh-binding site (AACTGGTT) was mutated to an EcoRI recognition site (AAGAATTC). *Drosophila* Schneider 2 (S2) cells were cultured in Schneider’s Media (Life Technologies) with 10% FBS (Omega Scientific) and 1% antibiotic/antimycotic (Life Technologies). Transient transfections were performed in technical triplicate with 900 ng reporter construct, 200 ng Grh-expression plasmid, and 100 ng of actin-renilla loading control plasmid using Effectene Transfection Reagent (Qiagen). Fold activation was calculated relative to luciferase controls transfected with 200 ng empty expression plasmid. Luciferase assays were performed on cell lysates using a Dual Luciferase Assay kit (Promega). Transient Grh expression was confirmed using a western blot with a previously validated antibody (Nevil et al., 2017).

### Formaldehyde-assisted isolation of regulatory elements

Formaldehyde-assisted isolation of regulatory elements (FAIRE) was performed as described in (McKay and Lieb, 2013). Briefly, wandering 3^rd^ instar larvae were dissected in 1xPBS such that the imaginal discs remained attached to the inverted cuticle. The prepared larvae were fixed in 4% formaldehyde for 10 minutes and subsequently the wing discs were further dissected and flash frozen to be stored at −80°C. Replicates of 40 wing discs each were perforated using bead beating 6x for 1 minute, with 2 minutes rest at 4°C. The chromatin was then sonicated using a Covaris S220 High Performance Ultrasonicator 4x for 30 seconds, 350 Watts power, 10% Duty factor, at 2°C. Phenol-chloroform extractions and ethanol precipitation were used to extract the accessible chromatin and purify the DNA. DNA concentrations were determined using a Qubit (ThermoFisher Scientific).

### Chromatin Immunoprecipitation

Chromatin Immunoprecipitation experiments were done with *lbl*^*GBE-*^ embryos and *w*^*1118*^ collected 11-12 hours after egg laying. Embryos were fixed using formaldehyde, chromatin extracted, and immunoprecipitations were performed using an αGrh antibody as described in (Li et al., 2008; Nevil et al., 2017). Enrichment was determined using biological triplicate in qPCR (described below).

### Quantitative PCR

DNA from either FAIRE or ChIP was prepared (as described above). Primers designed to span target regions (including controls) were used to perform qPCR in replicates using GoTaq qPCR Master Mix (Promega) (Table S1). Samples were analyzed in triplicate for each of three biological replicates and the fold change was calculated using fold enrichment over IgG (for ChIP) and the ΔΔC_t_ method (for FAIRE).

## Acknowledgements

We thank Marissa Gaskill and the Biochemistry Optical Core at the University of Wisconsin Madison for assistance in confocal imaging. We also thank the Bloomington Stock Center (NIH P40OD018537) and Dr. Sarah Bray for fly stocks.

## Data availability

The genomic data in this work was deposited in the Gene Expression Omnibus: accession number GSE137075. Strains and plasmids are available upon request.

## References

Abdusselamoglu, M. D., Eroglu, E., Burkard, T. R. and Knoblich, J. A. (2019). The transcription factor odd-paired regulates temporal identity in transit-amplifying neural progenitors via an incoherent feed-forward loop. Elife 8,.

Aleksic, J., Ferrero, E., Fischer, B., Shen, S. and Russell, S. (2013). The role of Dichaete in transcriptional regulation during Drosophila embryonic development. BMC Genomics 14, 861.

Attardi, L. D. and Tjian, R. (1993). Drosophila tissue-specific transcription factor NTF-1 contains a novel isoleucine-rich activation motif. Genes Dev. 7, 1341–1353.

Bellec, M., Radulescu, O. and Lagha, M. (2018). Remembering the past: Mitotic bookmarking in a developing embryo. Curr. Opin. Syst. Biol. 11, 41–49.

Benedyk, M. J., Mullen, J. R. and DiNardo, S. (1994). odd-paired: a zinc finger pair-rule protein required for the timely activation of engrailed and wingless in Drosophila embryos. Genes Dev. 8, 105–117.

Blythe, S. A. and Wieschaus, E. F. (2016). Establishment and maintenance of heritable chromatin structure during early Drosophila embryogenesis. Elife 5,.

Boglev, Y., Wilanowski, T., Caddy, J., Parekh, V., Auden, A., Darido, C., Hislop, N. R., Cangkrama, M., Ting, S. B. and Jane, S. M. (2011). The unique and cooperative roles of the Grainy head-like transcription factors in epidermal development reflect unexpected target gene specificity. Dev. Biol. 349, 512–522.

Bolger, A. M., Lohse, M. and Usadel, B. (2014). Trimmomatic: a flexible trimmer for Illumina sequence data. Bioinformatics 30, 2114–2120.

Bonn, S. and Furlong, E. E. M. (2008). cis-Regulatory networks during development: a view of Drosophila. Curr. Opin. Genet. Dev. 18, 513–520.

Bray, S. J. and Kafatos, F. C. (1991). Developmental function of Elf-1: an essential transcription factor during embryogenesis in Drosophila. Genes Dev. 5, 1672–83.

Bray, S. J., Johnson, W. A., Hirsh, J., Heberlein, U. and Tjian, R. (1988). A cis-acting element and associated binding factor required for CNS expression of the Drosophila melanogaster dopa decarboxylase gene. EMBO J. 7, 177–188.

Bray, S. J., Burke, B., Brown, N. H. and Hirsh, J. (1989). Embryonic expression pattern of a family of Drosophila proteins that interact with a central nervous system regulatory element. 3,.

Buecker, C., Srinivasan, R., Wu, Z., Calo, E., Acampora, D., Faial, T., Simeone, A., Tan, M., Swigut, T. and Wysocka, J. (2014). Reorganization of Enhancer Patterns in Transition from Naive to Primed Pluripotency. Cell Stem Cell 14, 838–853.

Buenrostro, J. D., Giresi, P. G., Zaba, L. C., Chang, H. Y. and Greenleaf, W. J. (2013). Transposition of native chromatin for fast and sensitive epigenomic profiling of open chromatin, DNA-binding proteins and nucleosome position. Nat. Methods 10, 1213–1218.

Caravaca, J. M., Donahue, G., Becker, J. S., He, X., Vinson, C. and Zaret, K. S. (2013). Bookmarking by specific and nonspecific binding of FoxA1 pioneer factor to mitotic chromosomes. Genes Dev. 27, 251–60.

Chen, W., Yi, J. K., Shimane, T., Mehrazarin, S., Lin, Y.-L., Shin, K.-H., Kim, R. H., Park, N.-H. and Kang, M. K. (2016). Grainyhead-like 2 regulates epithelial plasticity and stemness in oral cancer cells. Carcinogenesis 37, 500–510.

Chen, A. F., Liu, A. J., Krishnakumar, R., Freimer, J. W., DeVeale, B. and Blelloch, R. (2018). GRHL2-Dependent Enhancer Switching Maintains a Pluripotent Stem Cell Transcriptional Subnetwork after Exit from Naive Pluripotency. Cell Stem Cell 23, 226–238.e4.

Chou, T. B. and Perrimon, N. (1996). The autosomal FLP-DFS technique for generating germline mosaics in Drosophila melanogaster. Genetics 144, 1673–9.

Cirillo, L. A., Lin, F. R., Cuesta, I., Friedman, D., Jarnik, M. and Zaret, K. S. (2002). Opening of Compacted Chromatin by Early Developmental Transcription Factors HNF3 (FoxA) and GATA-4. Mol. Cell 9, 279–289.

Clark, E. and Akam, M. (2016). Odd-paired controls frequency doubling in Drosophila segmentation by altering the pair-rule gene regulatory network. Elife 5,.

Crémazy, F., Berta, P. and Girard, F. (2001). Genome-wide analysis of Sox genes in Drosophila melanogaster. Mech. Dev. 109, 371–375.

Davidson, E. H. (2006). The Regulatory Genome: Gene Regulatory Networks In Development And Evolution. Academic Press.

dos Santos, G., Schroeder, A. J., Goodman, J. L., Strelets, V. B., Crosby, M. A., Thurmond, J., Emmert, D. B., Gelbart, W. M. and FlyBase Consortium, the F. (2015). FlyBase: introduction of the Drosophila melanogaster Release 6 reference genome assembly and large-scale migration of genome annotations. Nucleic Acids Res. 43, D690–7.

Festuccia, N., Dubois, A., Vandormael-Pournin, S., Gallego Tejeda, E., Mouren, A., Bessonnard, S., Mueller, F., Proux, C., Cohen-Tannoudji, M. and Navarro, P. (2016). Mitotic binding of Esrrb marks key regulatory regions of the pluripotency network. Nat. Cell Biol. 18, 1139–1148.

Festuccia, N., Gonzalez, I., Owens, N. and Navarro, P. (2017). Mitotic bookmarking in development and stem cells. Development 144, 3633–3645.

Foo, S. M., Sun, Y., Lim, B., Ziukaite, R., O’Brien, K., Nien, C.-Y., Kirov, N., Shvartsman, S. Y. and Rushlow, C. A. (2014). Zelda potentiates morphogen activity by increasing chromatin accessibility. Curr. Biol. 24, 1341–1346.

Franco, H. L., Nagari, A. and Kraus, W. L. (2015). TNFα signaling exposes latent estrogen receptor binding sites to alter the breast cancer cell transcriptome. Mol. Cell 58, 21–34.

Gao, X., Vockley, C. M., Pauli, F., Newberry, K. M., Xue, Y., Randell, S. H., Reddy, T. E. and Hogan, B. L. M. (2013). Evidence for multiple roles for grainyhead-like 2 in the establishment and maintenance of human mucociliary airway epithelium.[corrected]. Proc. Natl. Acad. Sci. U. S. A. 110, 9356–61.

Garcia, M. and Stathopoulos, A. (2011). Lateral Gene Expression in Drosophila Early Embryos Is Supported by Grainyhead-Mediated Activation and Tiers of Dorsally-Localized Repression. PLoS One 6, e29172.

Gratz, S. J., Cummings, A. M., Nguyen, J. N., Hamm, D. C., Donohue, L. K., Harrison, M. M., Wildonger, J. and O’Connor-Giles, K. M. (2013). Genome Engineering of Drosophila with the CRISPR RNA-Guided Cas9 Nuclease. Genetics 194, 1029–1035.

Gross, D. S. and Garrard, W. T. (1988). Nuclease Hypersensitive Sites in Chromatin. Annu. Rev. Biochem. 57, 159–197.

Hamm, D. C., Larson, E. D., Nevil, M., Marshall, K. E., Bondra, E. R. and Harrison, M. M. (2017). A conserved maternal-specific repressive domain in Zelda revealed by Cas9-mediated mutagenesis in Drosophila melanogaster. PLOS Genet. 13, e1007120.

Harrison, M. M., Botchan, M. R. and Cline, T. W. (2010). Grainyhead and Zelda compete for binding to the promoters of the earliest-expressed Drosophila genes. Dev. Biol. 345, 248–255.

Harrison, M. M., Li, X.-Y., Kaplan, T., Botchan, M. R. and Eisen, M. B. (2011). Zelda Binding in the Early Drosophila melanogaster Embryo Marks Regions Subsequently Activated at the Maternal-to-Zygotic Transition. PLoS Genet. 7, e1002266.

Heinz, S., Benner, C., Spann, N., Bertolino, E., Lin, Y. C., Laslo, P., Cheng, J. X., Murre, C., Singh, H. and Glass, C. K. (2010). Simple combinations of lineage-determining transcription factors prime cis-regulatory elements required for macrophage and B cell identities. Mol. Cell 38, 576–89.

Hemphälä, J., Uv, A., Cantera, R., Bray, S. and Samakovlis, C. (2003). Grainy head controls apical membrane growth and tube elongation in response to Branchless/FGF signalling. Development 130, 249–58.

Huang, J. D., Dubnicoff, T., Liaw, G. J., Bai, Y., Valentine, S. A., Shirokawa, J. M., Lengyel, J. A. and Courey, A. J. (1995). Binding sites for transcription factor NTF-1/Elf-1 contribute to the ventral repression of decapentaplegic. Genes Dev. 9, 3177–3189.

Iwafuchi-Doi, M. (2019). The mechanistic basis for chromatin regulation by pioneer transcription factors. Wiley Interdiscip. Rev. Syst. Biol. Med. 11, e1427.

Iwafuchi-Doi, M. and Zaret, K. S. (2014). Pioneer transcription factors in cell reprogramming. Genes Dev. 28, 2679–2692.

Jacobs, J., Atkins, M., Davie, K., Imrichova, H., Romanelli, L., Christiaens, V., Hulselmans, G., Potier, D., Wouters, J., Taskiran, I. I., et al. (2018). The transcription factor Grainy head primes epithelial enhancers for spatiotemporal activation by displacing nucleosomes. Nat. Genet. 50, 1011–1020.

Kadauke, S., Udugama, M. I., Pawlicki, J. M., Achtman, J. C., Jain, D. P., Cheng, Y., Hardison, R. C. and Blobel, G. A. (2012). Tissue-Specific Mitotic Bookmarking by Hematopoietic Transcription Factor GATA1. Cell 150, 725–737.

Kaplan, T., Li, X.-Y., Sabo, P. J., Thomas, S., Stamatoyannopoulos, J. A., Biggin, M. D. and Eisen, M. B. (2011). Quantitative Models of the Mechanisms That Control Genome-Wide Patterns of Transcription Factor Binding during Early Drosophila Development. PLoS Genet. 7, e1001290.

Kent, W. J., Sugnet, C. W., Furey, T. S., Roskin, K. M., Pringle, T. H., Zahler, A. M. and Haussler, D. (2002). The human genome browser at UCSC. Genome Res. 12, 996–1006.

Kent, W. J., Zweig, A. S., Barber, G., Hinrichs, A. S. and Karolchik, D. (2010). BigWig and BigBed: enabling browsing of large distributed datasets. Bioinformatics 26, 2204–2207.

Kornberg, R. D. (1977). Structure of Chromatin. Annu. Rev. Biochem. 46, 931–954.

Kornberg, R. D. and Lorch, Y. (1999). Twenty-Five Years of the Nucleosome, Fundamental Particle of the Eukaryote Chromosome. Cell 98, 285–294.

Langmead, B. and Salzberg, S. L. (2012). Fast gapped-read alignment with Bowtie 2. Nat. Methods 9, 357–359.

Lee, H. and Adler, P. N. (2004). The grainy head transcription factor is essential for the function of the frizzled pathway in the Drosophila wing. Mech. Dev. 121, 37–49.

Li, X., MacArthur, S., Bourgon, R., Nix, D., Pollard, D. A., Iyer, V. N., Hechmer, A., Simirenko, L., Stapleton, M., Hendriks, C. L. L., et al. (2008). Transcription Factors Bind Thousands of Active and Inactive Regions in the Drosophila Blastoderm. PLoS Biol. 6, e27.

Li, H., Handsaker, B., Wysoker, A., Fennell, T., Ruan, J., Homer, N., Marth, G., Abecasis, G., Durbin, R. and 1000 Genome Project Data Processing Subgroup, 1000 Genome Project Data Processing (2009). The Sequence Alignment/Map format and SAMtools. Bioinformatics 25, 2078–9.

Li, X.-Y., Thomas, S., Sabo, P. J., Eisen, M. B., Stamatoyannopoulos, J. A. and Biggin, M. D. (2011). The role of chromatin accessibility in directing the widespread, overlapping patterns of Drosophila transcription factor binding. Genome Biol. 12, R34.

Liang, H.-L., Nien, C.-Y., Liu, H.-Y., Metzstein, M. M., Kirov, N. and Rushlow, C. (2008). The zinc-finger protein Zelda is a key activator of the early zygotic genome in Drosophila. Nature 456, 400–403.

Liao, Y., Smyth, G. K. and Shi, W. (2013). featureCounts: An efficient general-purpose program for assigning sequence reads to genomic features.

Liaw, G. J., Rudolph, K. M., Huang, J. D., Dubnicoff, T., Courey, A. J. and Lengyel, J. A. (1995). The torso response element binds GAGA and NTF-1/Elf-1, and regulates tailless by relief of repression. 9,.

Liu, Z. and Kraus, W. L. (2017). Catalytic-Independent Functions of PARP-1 Determine Sox2 Pioneer Activity at Intractable Genomic Loci. Mol. Cell 65, 589–603.e9.

Liu, Y., Pelham-Webb, B., Di Giammartino, D. C., Li, J., Kim, D., Kita, K., Saiz, N., Garg, V., Doane, A., Giannakakou, P., et al. (2017). Widespread Mitotic Bookmarking by Histone Marks and Transcription Factors in Pluripotent Stem Cells. Cell Rep. 19, 1283–1293.

Long, H. K., Prescott, S. L. and Wysocka, J. (2016). Ever-Changing Landscapes: Transcriptional Enhancers in Development and Evolution. Cell 167, 1170–1187.

Love, M. I., Huber, W. and Anders, S. (2014). Moderated estimation of fold change and dispersion for RNA-seq data with DESeq2. Genome Biol. 15, 550.

Luger, K., Dechassa, M. L. and Tremethick, D. J. (2012). New insights into nucleosome and chromatin structure: an ordered state or a disordered affair? Nat. Rev. Mol. Cell Biol. 13, 436–447.

Luo, Z., Gao, X., Lin, C., Smith, E. R., Marshall, S. A., Swanson, S. K., Florens, L., Washburn, M. P. and Shilatifard, A. (2015). Zic2 is an enhancer-binding factor required for embryonic stem cell specification. Mol. Cell 57, 685–694.

McDaniel, S. L., Gibson, T. J., Schulz, K. N., Fernandez Garcia, M., Nevil, M., Jain, S. U., Lewis, P. W., Zaret, K. S. and Harrison, M. M. (2019). Continued Activity of the Pioneer Factor Zelda Is Required to Drive Zygotic Genome Activation. Mol. Cell 74, 185–195.e4.

McKay, D. J. and Lieb, J. D. (2013). A common set of DNA regulatory elements shapes Drosophila appendages. Dev. Cell 27, 306–18.

Nambu, P. A. and Nambu, J. R. (1996). The Drosophila fish-hook gene encodes a HMG domain protein essential for segmentation and CNS development. Development 122, 3467–75.

Narasimha, M., Uv, A., Krejci, A., Brown, N. H. and Bray, S. J. (2008). Grainy head promotes expression of septate junction proteins and influences epithelial morphogenesis. J. Cell Sci. 121, 747–52.

Nevil, M., Bondra, E. R., Schulz, K. N., Kaplan, T. and Harrison, M. M. (2017). Stable Binding of the Conserved Transcription Factor Grainy Head to its Target Genes Throughout Drosophila melanogaster Development. Genetics 205, 605–620.

Nishino, H., Takano, S., Yoshitomi, H., Suzuki, K., Kagawa, S., Shimazaki, R., Shimizu, H., Furukawa, K., Miyazaki, M. and Ohtsuka, M. (2017). Grainyhead-like 2 (GRHL2) regulates epithelial plasticity in pancreatic cancer progression. Cancer Med. 6, 2686–2696.

Paré, A., Kim, M., Juarez, M. T., Brody, S. and McGinnis, W. (2012). The Functions of Grainy Head-Like Proteins in Animals and Fungi and the Evolution of Apical Extracellular Barriers. PLoS One 7, e36254.

Pédelacq, J.-D., Cabantous, S., Tran, T., Terwilliger, T. C. and Waldo, G. S. (2006). Engineering and characterization of a superfolder green fluorescent protein. Nat. Biotechnol. 24, 79–88.

Potier, D., Davie, K., Hulselmans, G., Naval Sanchez, M., Haagen, L., Huynh-Thu, V. A., Koldere, D., Celik, A., Geurts, P., Christiaens, V., et al. (2014). Mapping Gene Regulatory Networks in Drosophila Eye Development by Large-Scale Transcriptome Perturbations and Motif Inference. Cell Rep. 9, 2290–2303.

R Core Team (2017). R: A Language and Environment for Statistical Computing.

Raccaud, M. and Suter, D. M. (2018). Transcription factor retention on mitotic chromosomes: regulatory mechanisms and impact on cell fate decisions. FEBS Lett. 592, 878–887.

Raccaud, M., Friman, E. T., Alber, A. B., Agarwal, H., Deluz, C., Kuhn, T., Gebhardt, J. C. M. and Suter, D. M. (2019). Mitotic chromosome binding predicts transcription factor properties in interphase. Nat. Commun. 10, 487.

Raney, B. J., Dreszer, T. R., Barber, G. P., Clawson, H., Fujita, P. A., Wang, T., Nguyen, N., Paten, B., Zweig, A. S., Karolchik, D., et al. (2014). Track data hubs enable visualization of user-defined genome-wide annotations on the UCSC Genome Browser. Bioinformatics 30, 1003–1005.

Rifat, Y., Parekh, V., Wilanowski, T., Hislop, N. R., Auden, A., Ting, S. B., Cunningham, J. M. and Jane, S. M. (2010). Regional neural tube closure defined by the Grainy head-like transcription factors. Dev. Biol. 345, 237–245.

Russell, S. R., Sanchez-Soriano, N., Wright, C. R. and Ashburner, M. (1996). The Dichaete gene of Drosophila melanogaster encodes a SOX-domain protein required for embryonic segmentation. Development 122,.

Sandler, J. E. and Stathopoulos, A. (2016). Quantitative Single-Embryo Profile of Drosophila Genome Activation and the Dorsal-Ventral Patterning Network. Genetics 202, 1575–84.

Schindelin, J., Arganda-Carreras, I., Frise, E., Kaynig, V., Longair, M., Pietzsch, T., Preibisch, S., Rueden, C., Saalfeld, S., Schmid, B., et al. (2012). Fiji: an open-source platform for biological-image analysis. Nat. Methods 9, 676–682.

Schulz, K. N., Bondra, E. R., Moshe, A., Villalta, J. E., Lieb, J. D., Kaplan, T., McKay, D. J. and Harrison, M. M. (2015). Zelda is differentially required for chromatin accessibility, transcription factor binding, and gene expression in the early Drosophila embryo. Genome Res. 25, 1715–26.

Senga, K., Mostov, K. E., Mitaka, T., Miyajima, A. and Tanimizu, N. (2012). Grainyhead-like 2 regulates epithelial morphogenesis by establishing functional tight junctions through the organization of a molecular network among claudin3, claudin4, and Rab25. Mol. Biol. Cell 23, 2845–55.

Sharp, N. P. and Agrawal, A. F. (2012). Evidence for elevated mutation rates in low-quality genotypes. Proc. Natl. Acad. Sci. U. S. A. 109, 6142–6.

Sharp, N. P. and Agrawal, A. F. (2017). An experimental test of the mutation-selection balance model for the maintenance of genetic variance in fitness components. bioRxiv 193425.

Simon Andrews FastQC A Quality Control tool for High Throughput Sequence Data.

Soufi, A., Donahue, G. and Zaret, K. S. (2012). Facilitators and Impediments of the Pluripotency Reprogramming Factors’ Initial Engagement with the Genome. Cell 151, 994–1004.

Soufi, A., Garcia, M. F., Jaroszewicz, A., Osman, N., Pellegrini, M. and Zaret, K. S. (2015). Pioneer transcription factors target partial DNA motifs on nucleosomes to initiate reprogramming. Cell 161, 555–568.

Sun, Y., Nien, C.-Y., Chen, K., Liu, H.-Y., Johnston, J., Zeitlinger, J. and Rushlow, C. (2015). Zelda overcomes the high intrinsic nucleosome barrier at enhancers during Drosophila zygotic genome activation. Genome Res. 25, 1703–14.

Traylor-Knowles, N., Hansen, U., Dubuc, T. Q., Martindale, M. Q., Kaufman, L. and Finnerty, J. R. (2010). The evolutionary diversification of LSF and Grainyhead transcription factors preceded the radiation of basal animal lineages. BMC Evol. Biol. 10, 101.

Tremethick, D. J. (2007). Higher-Order Structures of Chromatin: The Elusive 30 nm Fiber. Cell 128, 651–654.

Uv, A. E., Thompson, C. R. and Bray, S. J. (1994). The Drosophila tissue-specific factor Grainyhead contains novel DNA-binding and dimerization domains which are conserved in the human protein CP2. Mol. Cell. Biol. 14, 4020–31.

Uv, A. E., Harrison, E. J. and Bray, S. J. (1997). Tissue-specific splicing and functions of the Drosophila transcription factor Grainyhead. 17,.

Venkatesan, K., McManus, H. R., Mello, C. C., Smith, T. F. and Hansen, U. (2003). Functional conservation between members of an ancient duplicated transcription factor family, LSF/Grainyhead. Nucleic Acids Res. 31, 4304–16.

Wang, S. and Samakovlis, C. (2012). Grainy Head and Its Target Genes in Epithelial Morphogenesis and Wound Healing. Academic Press.

Wickham, H. (2009). Ggplot2: elegant graphics for data analysis. Springer.

Wilanowski, T., Tuckfield, A., Cerruti, L., O’Connell, S., Saint, R., Parekh, V., Tao, J., Cunningham, J. M. and Jane, S. M. (2002). A highly conserved novel family of mammalian developmental transcription factors related to Drosophila grainyhead. Mech. Dev. 114, 37–50.

Wilczyński, B. and Furlong, E. E. M. (2010). Dynamic CRM occupancy reflects a temporal map of developmental progression. Mol. Syst. Biol. 6, 383.

Wittkopp, P. J. and Kalay, G. (2012). Cis-regulatory elements: molecular mechanisms and evolutionary processes underlying divergence. Nat. Rev. Genet. 13, 59–69.

Yao, L., Wang, S., Westholm, J. O., Dai, Q., Matsuda, R., Hosono, C., Bray, S., Lai, E. C. and Samakovlis, C. (2017). Genome-wide identification of Grainy head targets in Drosophila reveals regulatory interactions with the POU domain transcription factor Vvl. Development 144, 3145–3155.

Zaret, K. S. and Carroll, J. S. (2011). Pioneer transcription factors: establishing competence for gene expression. Genes Dev. 25, 2227–2241.

Zhang, Y., Liu, T., Meyer, C. A., Eeckhoute, J., Johnson, D. S., Bernstein, B. E., Nussbaum, C., Myers, R. M., Brown, M., Li, W., et al. (2008). Model-based Analysis of ChIP-Seq (MACS). Genome Biol. 9, R137.

Zinzen, R. P., Girardot, C., Gagneur, J., Braun, M. and Furlong, E. E. M. (2009). Combinatorial binding predicts spatio-temporal cis-regulatory activity. Nature 462, 65–70.

