## Supplemental Figures for "Establishment of chromatin accessibility by the conserved transcription factor Grainy head is developmentally regulated"

**Nevil Figure S1**

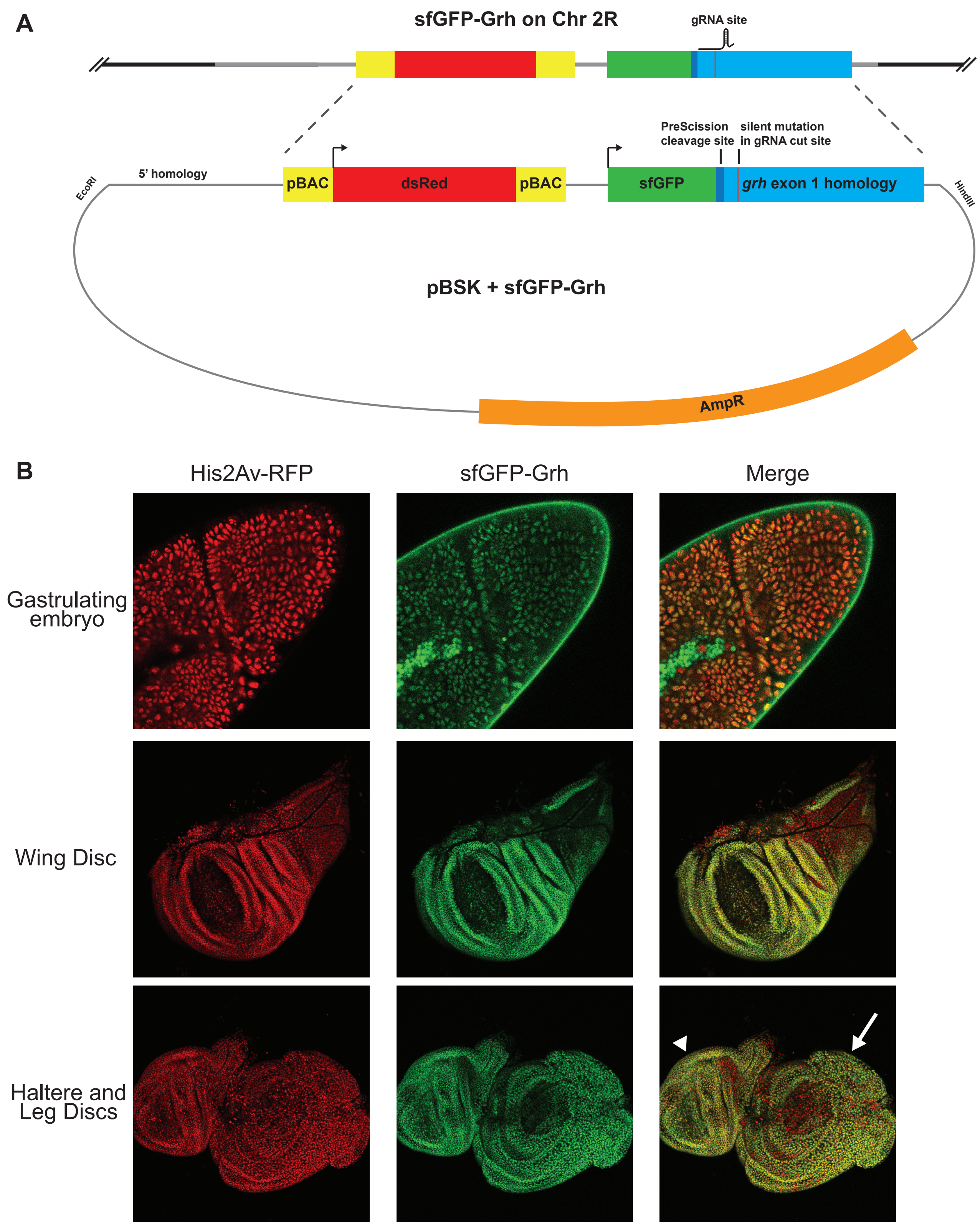


**Figure S1. Endogenous tagging of Grh with sfGFP.** (A) Scheme of sfGFP donor plasmid and predicted *grh* genomic locus after CRISPR/Cas9-mediated tagging. 1 kb homology arms (5' homology and exon 1 homology) flank a 3xP3-DsRed cassette inserted for screening. The DsRed cassette is flanked by piggyBac elements for precise excision after screening. sfGFP was inserted at the N-terminus of Grh, separated by a PreScission Protease cleavage site as a linker. (B) Confocal microscopy images of a *sfGFP-Grh; His2Av-RFP* embryo at gastrulation, larval wing disc, haltere disc (arrow head), and leg disc (arrow).

### Nevil Figure S2

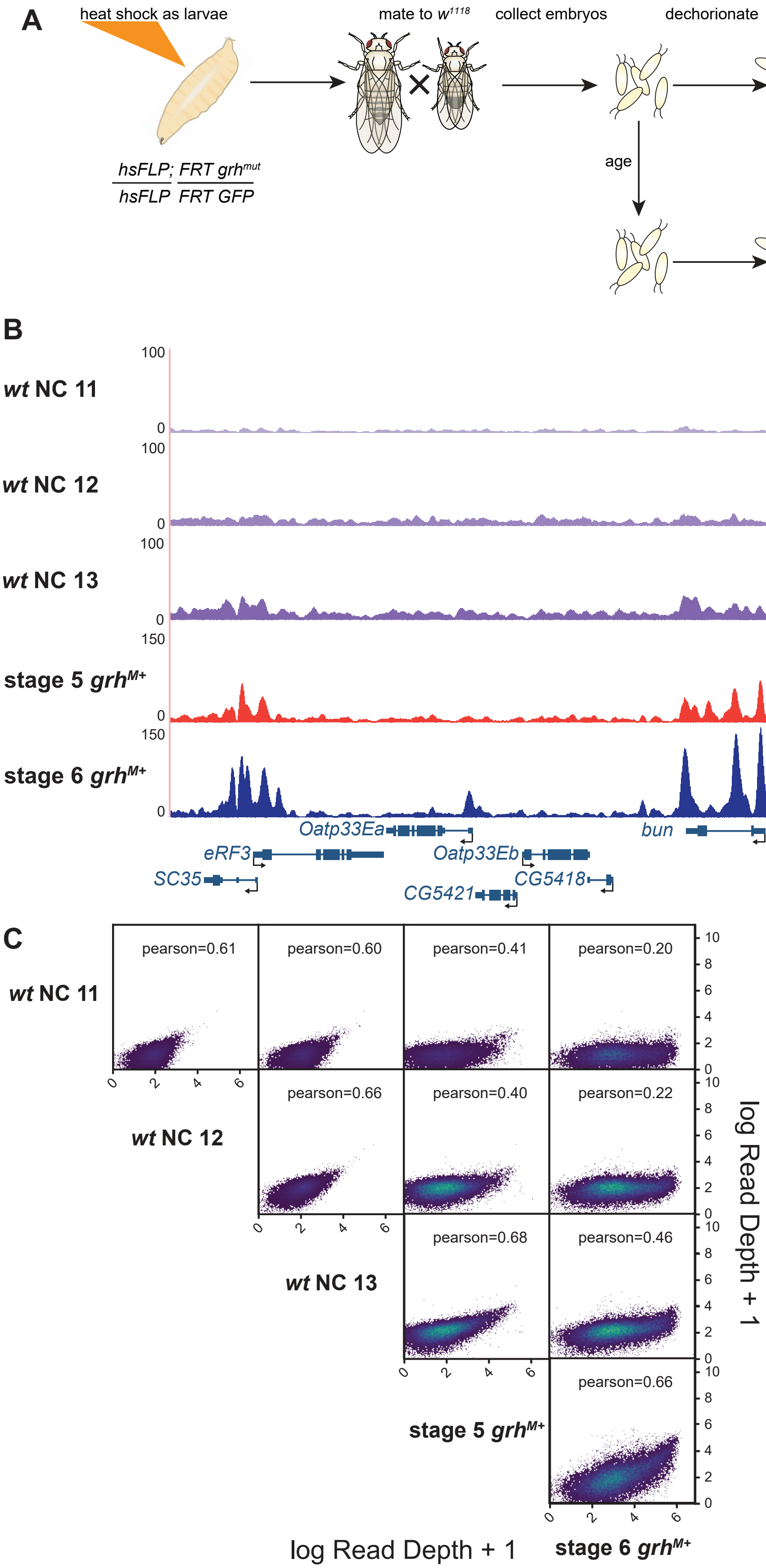


**Figure S2. ATAC-seq on embryos depleted of maternally encoded Grh.** (A) Scheme of heat shock FLP-FRT protocol used to generate germline mitotic clones lacking maternally encoded Grh. (B) UCSC genome browser track with published nuclear cycle (NC) 11, 12, and 13 ATAC-seq data from wild-type embryos (Blythe and Wieschaus, 2016) and wild-type stage 5 and stage 6 embryos. (C) Pearson correlation plots of published nuclear cycle (NC) 11, 12, and 13 ATAC-seq data from wild-type embryos (Blythe and Wieschaus, 2016) and wild-type stage 5 and stage 6 embryos at accessible regions of chromatin. Higher Pearson correlation coefficients indicate greater similarity between datasets. Data are transformed using log read depth + 1 for plotting, not for correlation calculation.

**Nevil Figure S3**

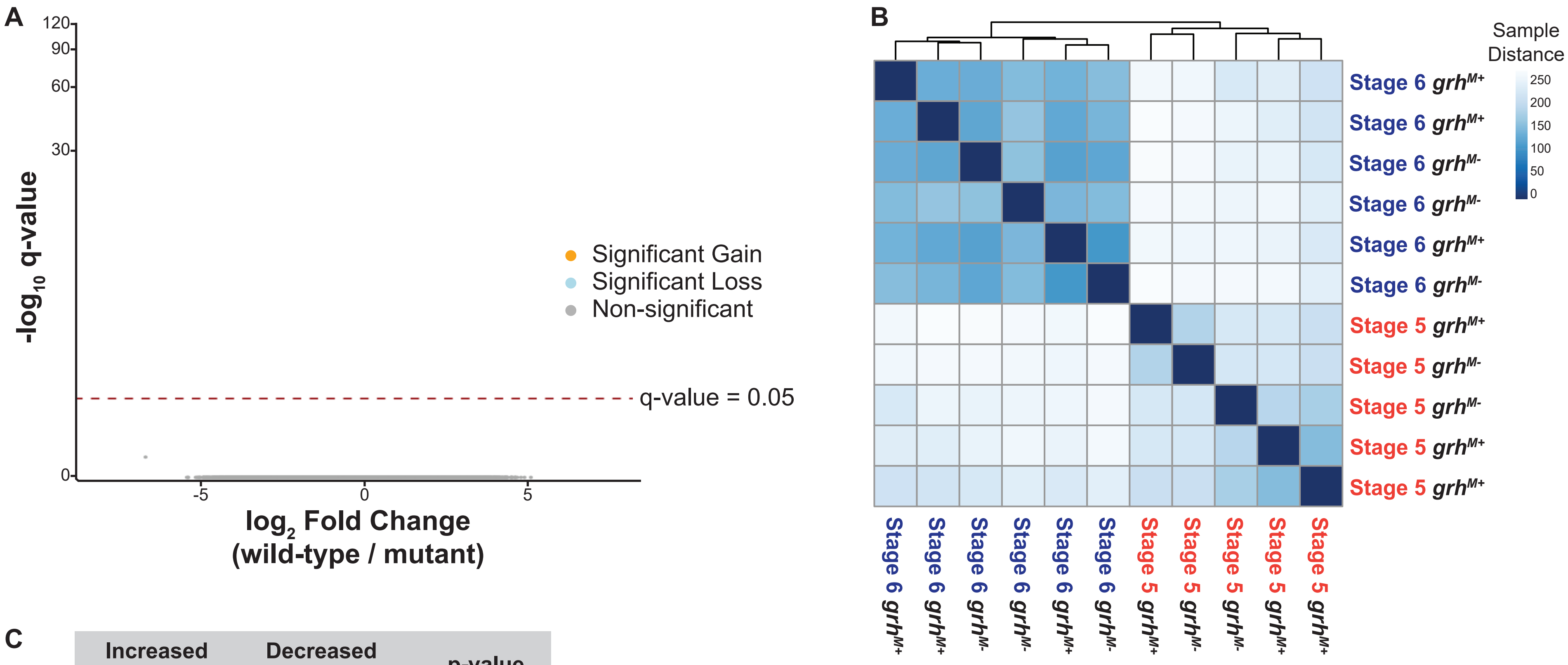


C

| | Increased Accessibility | Decreased Accessibility | p-value |
|---|---|---|---|
| **Grh** | **13.9%** | 4.5% | $< 2.2 \times 10^{-22}$ |
| **Zld** | 22.5% | **33.8%** | $< 2.2 \times 10^{-22}$ |

**Figure S3. Stage is a better predictor of clustering than genotype.** (A) Volcano plots of all accessible regions identified using the DESeq2 package in comparisons between stage 6 *grh* maternal depletion and heterozygous siblings (Love et al., 2014). Significance of change in accessibility reported by $-\log_{10}$ q-value on the y-axis, and magnitude of change by $\log_2$ fold change on the x-axis. Regions that significantly gain (orange) or lose (blue) accessibility are defined as those with a q-value < 0.05. Non-significant changes are those with a q-value > 0.05 (gray). (B) Hierarchical clustering of sample distances for single embryo ATAC-seq replicates of stage 5 and stage 6 maternal depletions ($grh^{M-}$) and sibling controls ($grh^{M+}$). Color and phylogenetic tree indicate sample-to-sample relatedness. (C) Percentages of ChIP-seq Grh (2-3hr AEL embryos) and Zld (cycle 14 embryos) peaks that overlap with sites that significantly increase or decrease in chromatin accessibility during gastrulation. p-values calculated using a two-sided Fisher's exact test comparing the enrichment of each transcription factor at differentially accessible sites.

Nevil Figure S4

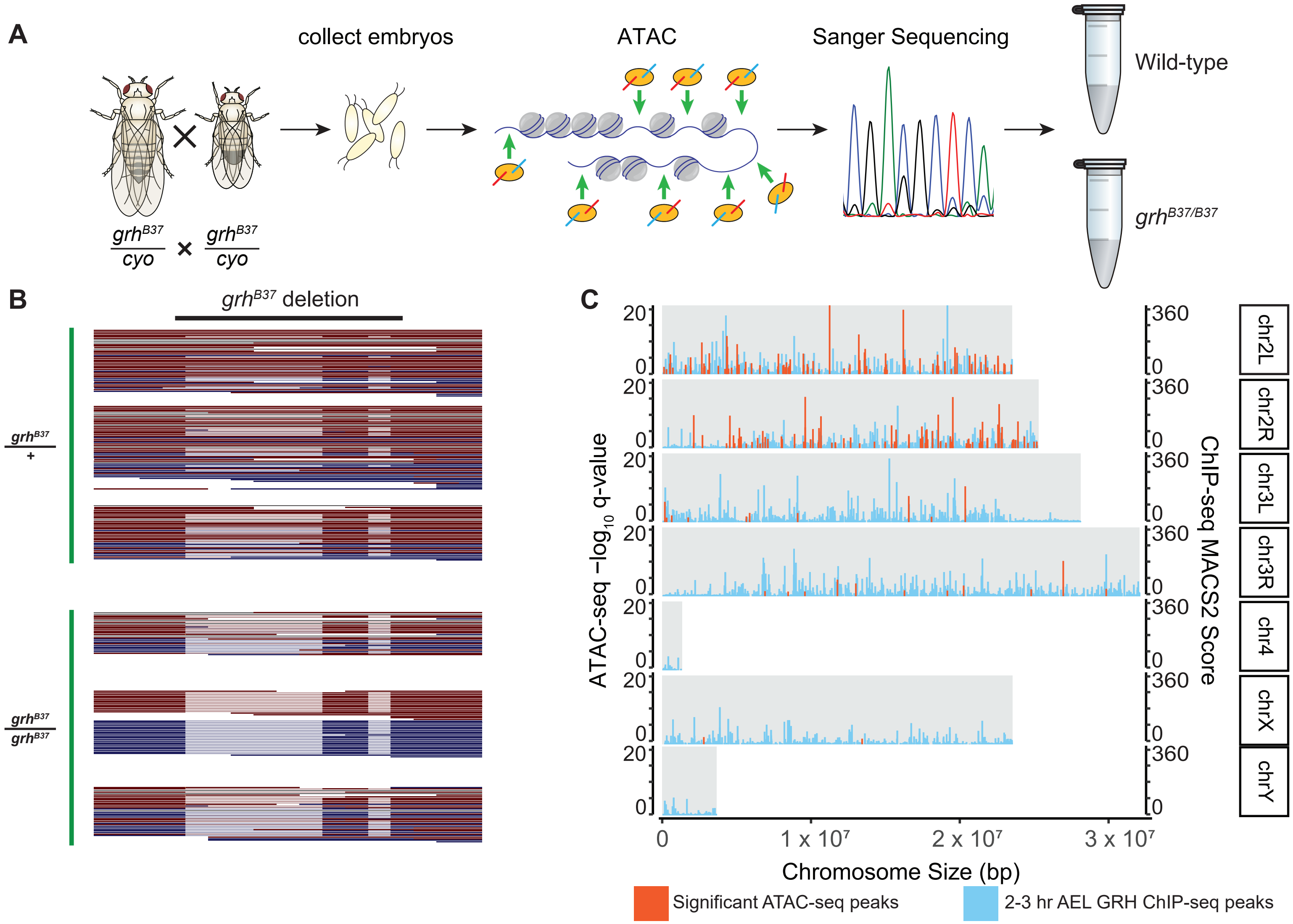


**Figure S4. ATAC-seq on embryos depleted of zygotically expressed Grh.** (A) Scheme used to collect and identify *grh*$^{B37/B37}$ embryos. Single embryos were collected for ATAC. After library preparation, the region spanning the *grh*$^{B37}$ deletion was amplified by PCR and sequenced. Homozygous mutants were selected. (B) ATAC-seq reads aligned (red = reverse strand, blue = forward strand) to the wild type sequence that is deleted in the *grh*$^{B37}$ allele. Each line is an individual read that spans the deletion, each block of reads represents a single replicate (three per genotype). Thick lines indicate overlap to the reference (wild type) sequence, whereas thin lines indicate lack of sequence overlap from the mutant allele. The lack of wild type, spanning reads in the homozygous mutant indicate correct identification of mutants. (C) Location of significantly changing regions (red bars) of accessibility show the largest number and most significant (-log$_{10}$ q-value) are located on chromosome 2, the chromosome where the *grh*$^{B37}$ allele has been maintained over a balancer allowing mutations to accrue and limiting mapping of sequencing reads. Plotting the location of Grh ChIP-seq peaks (blue bars, height by MACS2 score column) shows that Grh binding is not restricted to chromosome 2.

### Nevil Figure S5

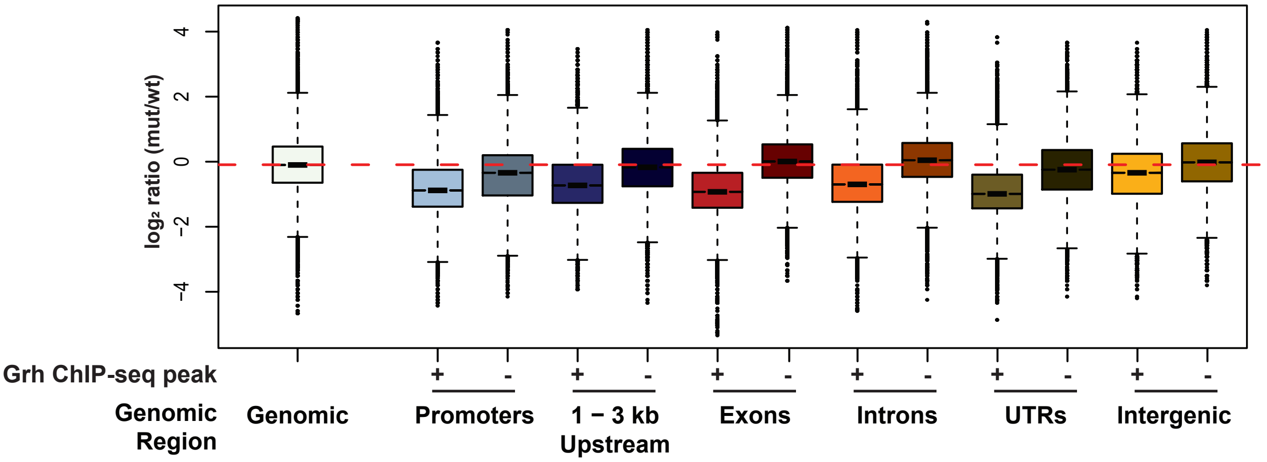


**Figure S5. Genomic regions that overlap with a Grh peak lose chromatin accessibility in the absence of Grh.** Changes in chromatin accessibility were calculated by taking random 100 bp windows within the indicated genomic features that either contained a Grh-binding site or did not (Nevil et al., 2017). The $\log_2$ ratio (*grh* mutant/ wild-type) of the read depth normalized for depth and breadth was calculated for each. The "Genomic" group indicates the signal from random genomic regions which are largely free of Grh binding.

**Nevil Figure S6**

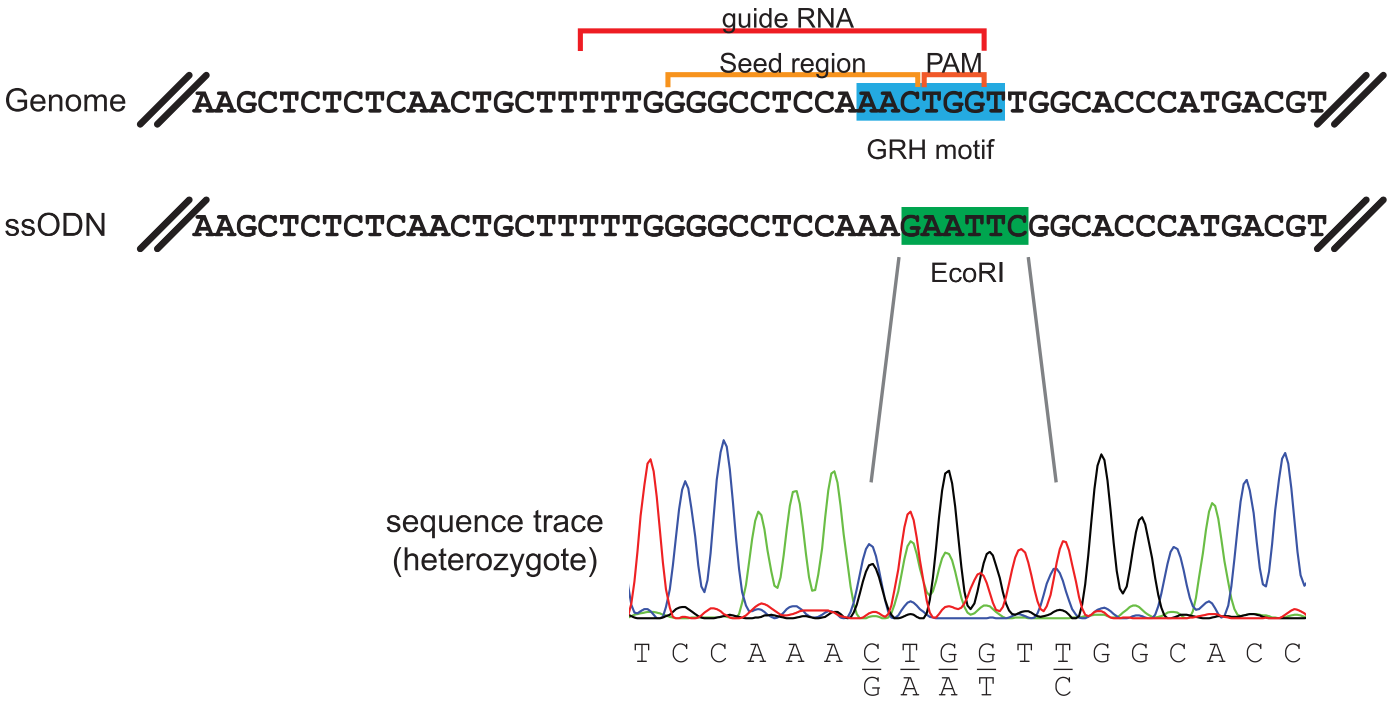


**Figure S6. ssODN design and mutation of Grh-binding site in the lbl promoter.** A 135-nucleotide single stranded oligodeoxynucleotide (ssODN) donor template was designed across the *lbl* promoter to mutate the canonical Grh motif. In the wild-type genomic locus the Grh motif overlaps the guideRNA-recognition site. Successful mutation to an EcoRI site sequence abrogates guideRNA recognition by mutating both the seed region and PAM. Mutants lines were identified and sequenced.

Nevil Figure S7

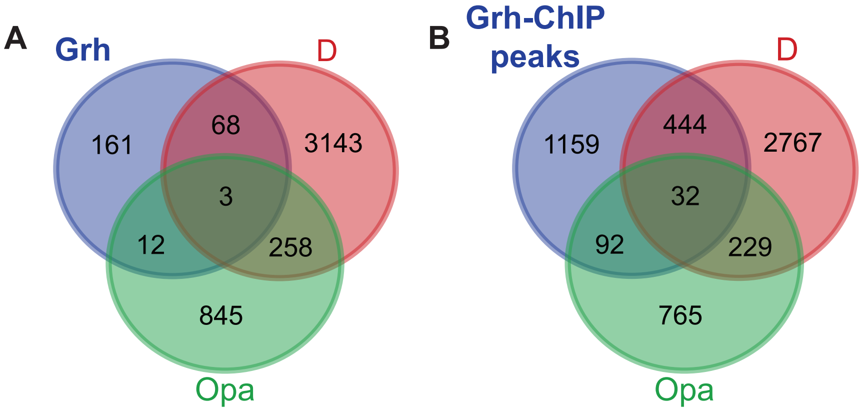


**Figure S7. Overlap of Grh, Opa and D binding motifs in regions that increase in chromatin accessibility during gastrulation.** (A) Overlap of regions that gain accessibility during gastrulation and contain motifs (as defined by HOMER) for the indicated factors. (B) Overlap of regions that gain accessibility during gastrulation and have a Grh-binding site as defined by ChIP-seq and contain motifs (as defined by HOMER) for Opa or D.

**Nevil Table S1**

| Name | Sequence |
|---|---|
| SrF | AGAGCAACGAAACTCACCAC |
| SrR | CGTCATATGTCTCGAATGTATGGT |
| SloF | AGCCCATCTAGGGTGAAGTA |
| SloR | TGATGCGGGTAAGTCAACTG |
| LblF | CTCTTTCTTGGCCAAGCTCTCT |
| LblR | GAGGCGGCGTTTCCAAT |
| Ash2F | TCGAAGCAAGAGCCAGAG |
| Ash2R | GTCGCCAAAGAGCGATAAAC |
| closed1F | TGTAGTTTGGGTCGCAGC |
| closed2R | AAAGGCAGTTGAGAATTGTTCG |
| closed1F | TTGGCAATTGCTACGGATTT |
| closed2R | TATGTGATTATGGCCGACGA |

**Table S1. Primers Used for qPCR**
